# Dynamics of the Cortico-Cerebellar Loop Fine-Tune Dexterous Movement

**DOI:** 10.1101/637447

**Authors:** Jian-Zhong Guo, Britton Sauerbrei, Jeremy D. Cohen, Matteo Mischiati, Austin Graves, Ferruccio Pisanello, Kristin Branson, Adam W. Hantman

**Author notes:** These authors contributed equally (Jian-Zhong Guo, Britton Sauerbrei). These authors contributed equally (Jeremy D. Cohen, Matteo Mischiati).

## Abstract

Skillful control of movement requires coordination between brain areas that are reciprocally connected through polysynaptic pathways, forming closed loops. A prominent loop in mammals runs between cerebral cortex and cerebellum, which individually contribute to skilled arm control. But how and why do these regions interact? Here, we studied the mouse cortico-cerebellar loop by optogenetically perturbing the pontine nuclei (PN), which receive direct cortical inputs and project only to cerebellum. PN stimulation during rest propagated into cerebellar cortex, but the effect of stimulation was transformed downstream into a wide range of patterns in the deep cerebellar nuclei (DCN) and reduced to transient excitation in motor cortex. PN stimulation in a cued reaching task altered arm kinematics and impaired performance. Cerebellar and cortical dynamics during movement were not dominated by PN stimulation, but altered in line with behavioral changes. These results suggest that the cortico-cerebellar loop fine-tunes motor commands during skilled reaching.

## Introduction

Motor cortex and cerebellum are critical for the control of voluntary arm movements. In primates, neurons in both regions are tuned to reach direction (Fortier et al., 1989; Georgopoulos et al., 1982), though cerebellar neurons exhibit more phasic firing patterns, weaker tuning, less holding-related activity, and higher trial-to-trial variability (Fortier et al., 1993). In mice, neural activity in motor cortex and cerebellum is modulated during reaching (Becker and Person, 2019; Sauerbrei et al., 2020). Experimental perturbations and lesions of motor cortex block movement initiation or disrupt reach kinematics in animal models (Fogassi et al., 2001; Galiñanes et al., 2018; Guo et al., 2015; Lawrence and Kuypers, 1968; Miri et al., 2017; Passingham et al., 1983; Sauerbrei et al., 2020), and stroke and neurological disease affecting arm motor cortex lead to weakness or paralysis of the arm in human patients (Krakauer and Carmichael, 2017). Cerebellar damage, disease, or perturbation results in timing deficits, ataxia, tremor, and dysmetria of voluntary arm movements, including reaching and grasping (Becker and Person, 2019; Dow and Moruzzi, 1958; Klockgether, 2000; Nashef et al., 2019).

Anatomically, motor cortex and cerebellum are reciprocally connected, forming the cortico-cerebellar loop (Fig. 1A-B). Cerebral cortex projects to the cerebellum through its monosynaptic projections to the pontine nuclei (PN), which include the basal pons and the reticulotegmental nucleus (Brodal and Bjaalie, 1992; Brodal and Jansen, 1946; Cajal, 1898; Leergaard et al., 2004; Legg et al., 1989; Mihailoff et al., 1985). The PN project to the cerebellum, their sole output target, providing mossy fiber inputs to the cerebellar cortex as well as collaterals to the deep cerebellar nuclei (DCN), the output stage of the cerebellum. The major synaptic partners of PN mossy fibers are Golgi cells and granule cells. Granule cells excite Purkinje cells, which in turn inhibit the DCN. The DCN projects to the motor thalamus, which projects back to motor cortex, closing the loop.

**Figure 1:**
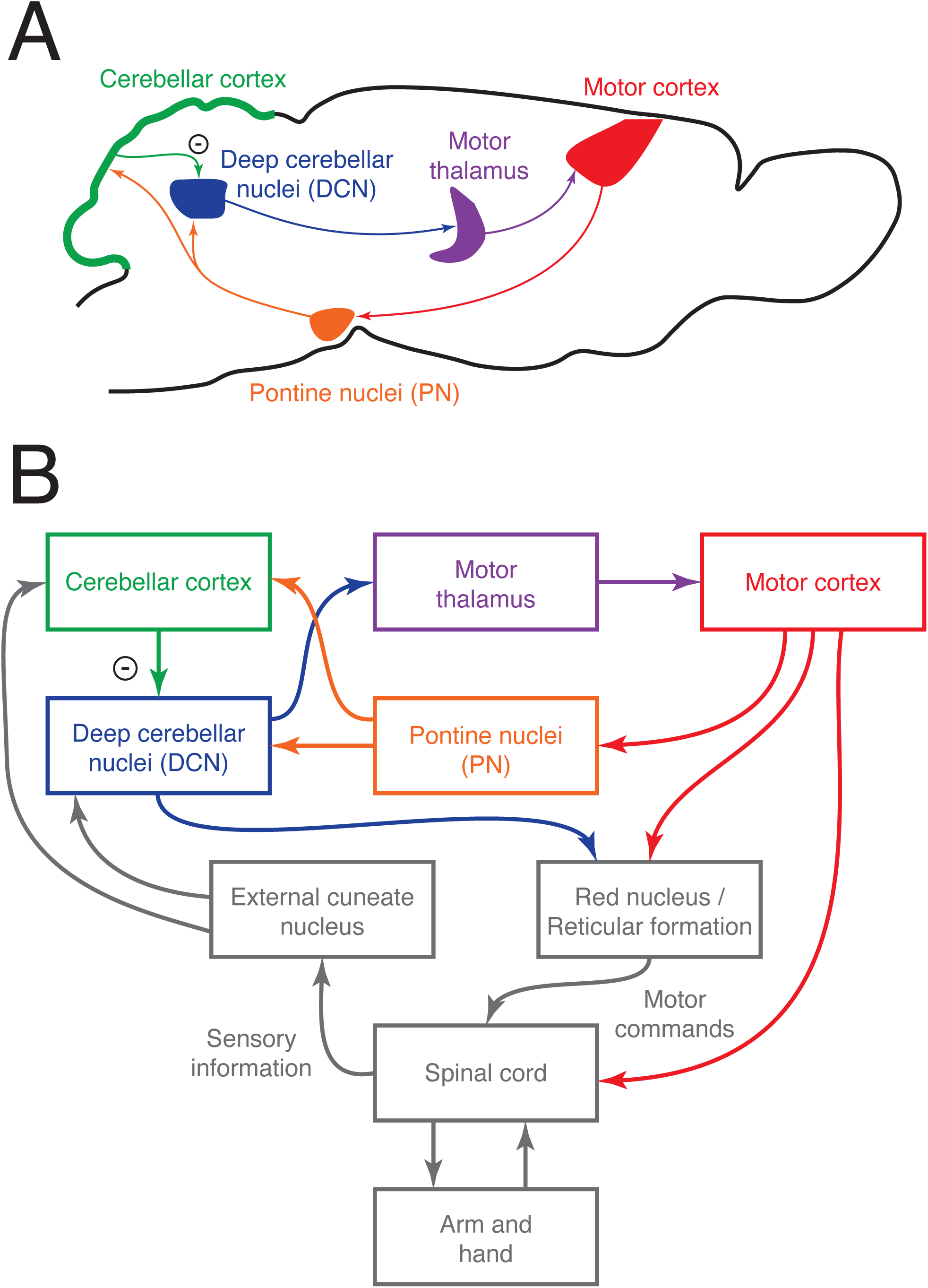
The cortico-cerebellar loop for arm control in the mouse. (A) Schematic of the main anatomical connections within the loop. (B) Relationship between the cortico-cerebellar loop and the sensorimotor periphery.

Several studies have investigated the cerebello-cortical portion of this loop. In primates, disruption of cerebellar output by cooling or high-frequency microstimulation impairs upper limb movement and alters motor cortical activity (Brooks et al., 1973; Meyer-Lohmann et al., 1975, 1977; Nashef et al., 2019). During the planning of tongue movements, a cerebellar perturbation disrupts cortical activity, a cortical perturbation disrupts cerebellar activity, and both disrupt motor planning (Gao et al., 2018). Thus, experimental manipulation of the DCN has provided some evidence that the cerebellum influences cortical activity related to movement. However, it remains unclear how selective perturbation of cortico-cerebellar projections impacts the rest of the loop and behavior.

The unique anatomical attributes of the PN make them a near ideal experimental target for testing the function of the descending cortico-cerebellar projections. In primates, PN neurons are modulated during arm and eye movements, and are tuned to movement direction (Matsunami, 1987; Tziridis et al., 2009). Although damage to the PN in humans results in motor impairments (Schmahmann et al., 2004), selective PN lesions in animal models have been difficult, as such lesions tend to also damage the corticospinal and medial lemniscus tracts, which run directly through the middle of the nuclei. Nonetheless, the effects of PN lesions have been studied, and the most prominent impairments have involved visual sensorimotor behaviors (Levesque et al., 1986; Stein and Glickstein, 1992) and gap crossing (Jenkinson and Glickstein, 2000). Optogenetic PN inhibition was recently shown to impact cortico-cerebellar coupling during arm movement (Wagner et al., 2019), but effects on behavior were limited to an increase in movement duration when the perturbation was applied repeatedly. Here, we use optogenetic excitation to examine how PN output influences movement through its effects on downstream activity in the DCN and cortex. We find that cortico-cerebellar dynamics fine-tune reach kinematics, enabling skilled motor performance.

## Results

### Firing properties of PN neurons during a dexterous, cortically-dependent behavior

Although the PN are the major conduit between cortex and cerebellum, little is known about the activity of PN neurons during behavior in mice that require these structures. We used a reach-to-grasp task (Fig. 2A, upper left and lower), which previous work has shown to depend on motor cortex (Guo et al., 2015; Sauerbrei et al., 2020) and cerebellum (Becker and Person, 2019). Because the regions of the PN receiving motor cortical input are small and deep in the brain, we targeted our recordings using a combination of high-density electrophysiology and optogenetic stimulation of corticopontine fibers (Fig. 2A, upper right). We performed the experiments in Sim1-Cre X Ai32 mice, which express ChR2 in layer 5 pyramidal tract neurons projecting to the PN (Gerfen et al., 2013). During the recording session, we slowly lowered a 960-site, 384-channel Neuropixels probe coated in fluorescent dye towards the PN (Fig. 2B). As the probe approached its target, we stimulated forelimb motor cortex with an optical fiber coupled to a 473 nm laser using a one-second, 10 Hz pulse train. This resulted in bursts of spiking activity in the target zones, verifying that the probe was in a region of the pontine nuclei receiving motor cortical input (Fig. 2C).

**Figure 2:**
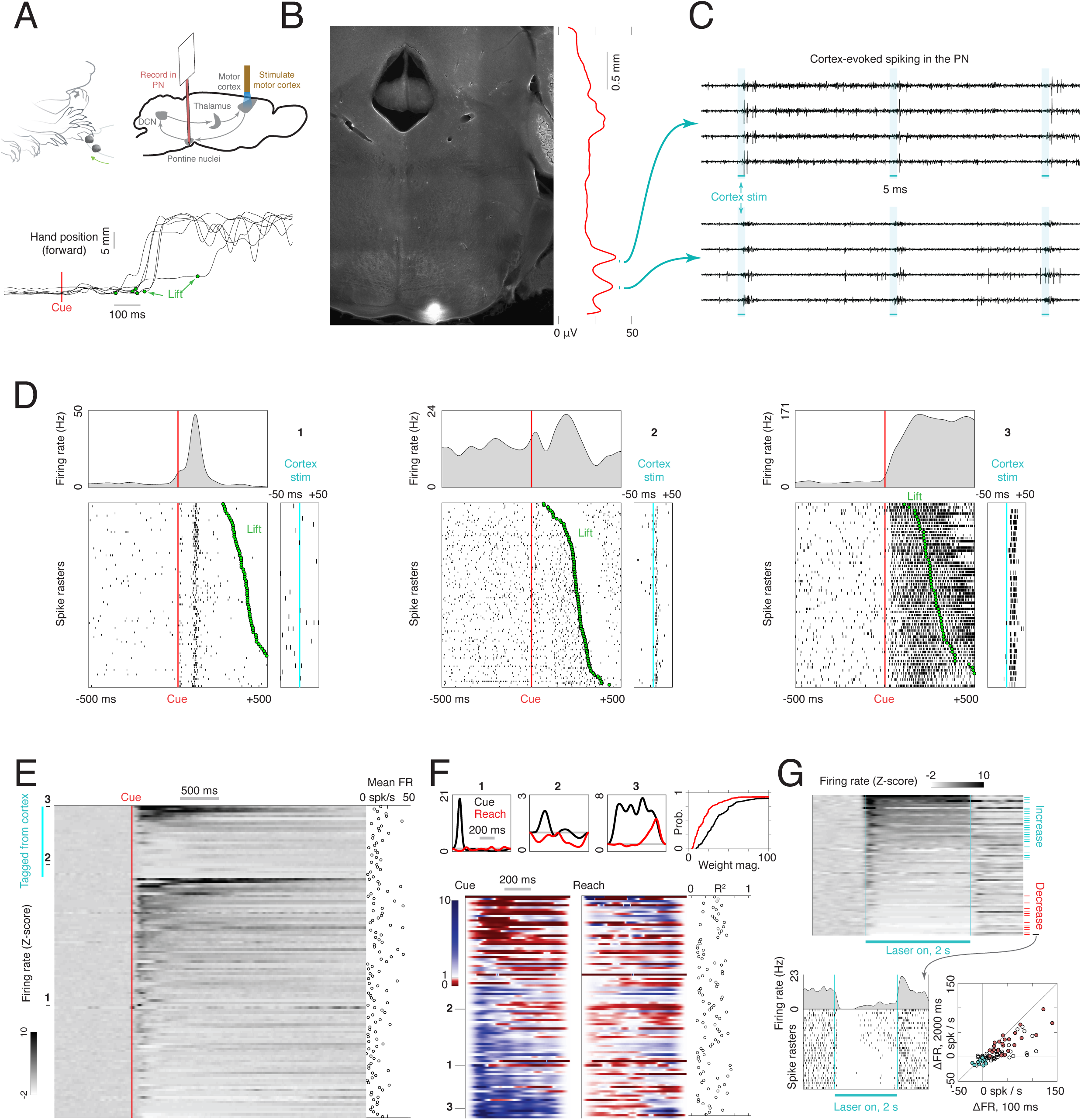
Firing properties of PN neurons during a cortically-dependent reach-to-grab task. (A) Experimental setup and reach kinematics. Upper left: mice were head-fixed and trained to reach to and grab a pellet of food following an acoustic cue. Upper right: Strategy for recording from motor-cortex-recipient neurons in the pontine nuclei. An optical fiber was placed over the forelimb motor cortex of mice expressing ChR2 in pyramidal tract neurons (Sim1-Cre X Ai32). A 384-channel Neuropixels probe was lowered into the pontine nuclei as a laser pulse train was delivered to cortex. After these stimuli were delivered, the animal performed the task as PN neurons were recorded. Lower: example hand trajectories aligned to the cue. The green dots indicate lift, when the animal initiates the reach-to-grab sequence. (B) Histological section showing the tip of the probe in the PN. Red line shows the amplitude of multi-unit activity on the probe in response to cortical stimulation as a function of depth. (C) Raw data from eight channels near the bottom of the probe showing activity evoked in the PN by motor cortex stimulation. (D) Spike rasters and firing rates for example neurons in the task. The timing of lift is indicated by the green dots. The panels on the right show each unit’s response to cortical stimulation. (E) Firing rate Z-scores for all PN neurons (n = 129), sorted according to whether the cells responded to cortical stimulation and by the magnitude of the response (n = 30 neurons were responsive). Inset at the right shows mean firing rates. (F) Effect of the cue and reaching on the firing rates of PN neurons (see Methods). Upper: cue (black) and reach (red) components from generalized linear models (GLMs) for the three example neurons in (D). Gray horizontal lines at the value of 1 correspond to no effect on firing rate; values less than one correspond to a decrease, and values greater than 1 correspond to an increase. Lower: cue and reach components for all neurons with R^2^ > .1 (n = 103). Blue values indicate firing rate increases, and red values indicate decreases. Inset at right shows R^2^ for each neuron. (G) Response of PN neurons to a two-second, 40 Hz stimulation of motor cortex. Upper: firing rate Z-scores for each neuron, aligned to laser onset. Blue ticks indicate neurons with a sustained increase, and red ticks indicate neurons with a decrease. Lower left: laser-aligned spike rasters and firing rate for an example neuron suppressed by motor cortical stimulation. Lower right: scatterplot of the change in firing rate from baseline over the entire two-second stimulation window vs over the initial 100 ms of stimulation. Color coding as in the upper panel.

We recorded the activity of 129 PN neurons, which exhibited a wide variety of patterns during the task (Fig. 2D-E, Supplemental Fig. 1). A subset of these neurons (n = 30) received input from motor cortex, revealed by short-latency responses to optogenetic stimulation of motor cortex (right inset of neurons 2 and 3 in Fig. 2D). In order to quantify the response of PN neurons to the cue and movement onset, we fit a generalized linear model (GLM) for each neuron (see Methods). This produced two curves, one indicating how the cue influenced firing rate over time, and the other indicating the influence of movement onset. These curves are shown for three example neurons in Fig. 2F (upper); these are the same neurons shown in Fig. 2D. Neuron 1 exhibits a sharp, transient increase in firing rate following the cue, but little change around lift. Neuron 2 has an increase following the cue, but a decrease following lift. Neuron 3 shows a rapid and sustained increase following cue, and a smaller, delayed increase following lift. The cue and reach curves are shown for all neurons for which we were able to fit a GLM (defined as those cells for which the R^2^ from the regression of observed firing rates on the firing rates predicted by the GLM exceeded .1, n = 103/129) in Fig. 2F (lower). Some cells were modulated around movement onset, consistent with previous results in primates (Matsunami, 1987; Tziridis et al., 2009), but the GLM weights for cue were typically larger than the weights for reach (Fig. 2F, upper right; p = 2.2e-9, signed rank test). Some previous work has suggested that different modalities of information remain segregated in the pons (Mihailoff et al., 1985; Schwarz and Thier, 1995; Wiesendanger and Wiesendanger, 1982), consistent with the similarity in neural dynamics in motor cortex and cerebellar granule cells (Wagner et al., 2019). Others, however, have suggested that PN neurons may integrate inputs from multiple cortical regions (Potter et al., 1978; Rüegg and Wiesendanger, 1975). Our observation that the activity of some motor cortex-recipient PN neurons is aligned both to the cue and movement suggests that these neurons might integrate signals of multiple modalities.

In rodents, there appear to be very few local inhibitory neurons within the pontine nuclei (Brodal et al., 1988). Inhibitory inputs to the PN have been identified in the zona incerta, anterior pretectal nucleus, and reticular formation (Border et al., 1986), but it is unclear whether these circuits can be recruited by descending cortical signals to suppress PN spiking. In order to identify possible cortically-driven feedforward inhibition onto PN neurons, we delivered a two-second, 40 Hz sinusoidal laser stimulus to cortex. Some neurons (n = 26) maintained elevated firing rates during this long stimulation, but others showed a sustained decrease (n = 12; Fig. 2G, upper, lower left). In some cases, these decreases were not preceded by transient excitation; this suggests that these neurons receive feedforward inhibition (Fig. 2G, lower right). Thus, motor cortex may be capable not only of exciting PN neurons through direct pyramidal tract projections, but also suppressing them through polysynaptic inhibitory routes. This might enable cortical output to the cerebellum to be gated off during motor preparation (Kaufman et al., 2014) or the execution of non-cortically-dependent behaviors (Miri et al., 2017).

### Propagation of PN signals across the cortico-cerebellar loop

How are PN signals transformed across the cortico-cerebellar loop? We addressed this question by optogenetically stimulating PN neurons and recording neural activity across the loop. Using the Slc17a7-Cre mouse line we generated (Huang et al., 2013) that expresses Cre recombinase selectively in PN neurons, we drove expression of ChR2 by injecting AAV2/1-FLEX-rev-ChR2-tdTomato into these mice. A tapered optical fiber was implanted in the PN (Fig. 3A, left), exploiting its sub-micrometer implant cross section and the smooth profile of the fiber, which make the targeted region easier to reach with respect to standard flat-cleaved fiber optics (Pisanello et al., 2017). We stimulated PN neurons with 473 nm light at 8-20 mW for 2 s at 40 Hz, a frequency near the upper limit for driving ChR2 (Berndt et al., 2011). We characterized the postsynaptic effects of stimulation by recording in the cerebellar cortex with a high-density Neuropixels probe. PN stimulation entrained multi-unit activity in the cerebellar cortex at 40 Hz (Fig. 3A-C). This multi-unit activity likely reflects combined signals from mossy fiber terminals, granule cells, Purkinje cells, and interneurons. Next, we examined how the effects of PN stimulation propagated into the deep cerebellar nuclei (DCN; Fig. 3D), which receives input from inhibitory Purkinje cells of the cerebellar cortex, as well as collateral excitatory inputs from the PN. We targeted 64-channel silicon probes to the anterior interpositus nucleus during optogenetic stimulation of the PN. We selected this portion of the DCN because it receives PN input, is known to contain arm-related neurons, is easily targeted for recording, and sends ascending projections to motor thalamus. DCN neurons (n = 139) exhibited a wide range of laser-aligned responses, including sustained or transient excitation or inhibition, slow ramps in activity, and combinations of these (Fig. 3D-E). Does the modulation of cerebellar output by PN stimulation propagate into motor cortex? To address this question, we again activated PN neurons with ChR2 and recorded neural activity in layer 5 of motor cortex (Fig. 4A). PN stimulation for 2 s evoked neural responses in 116/848 motor cortical neurons, but unlike the responses in the DCN, these were stereotypical, mostly consisting of transient excitation (Fig. 4B, left). These data show how PN signals can be transformed as they propagate across the entire cortico-cerebellar loop.

**Figure 3:**
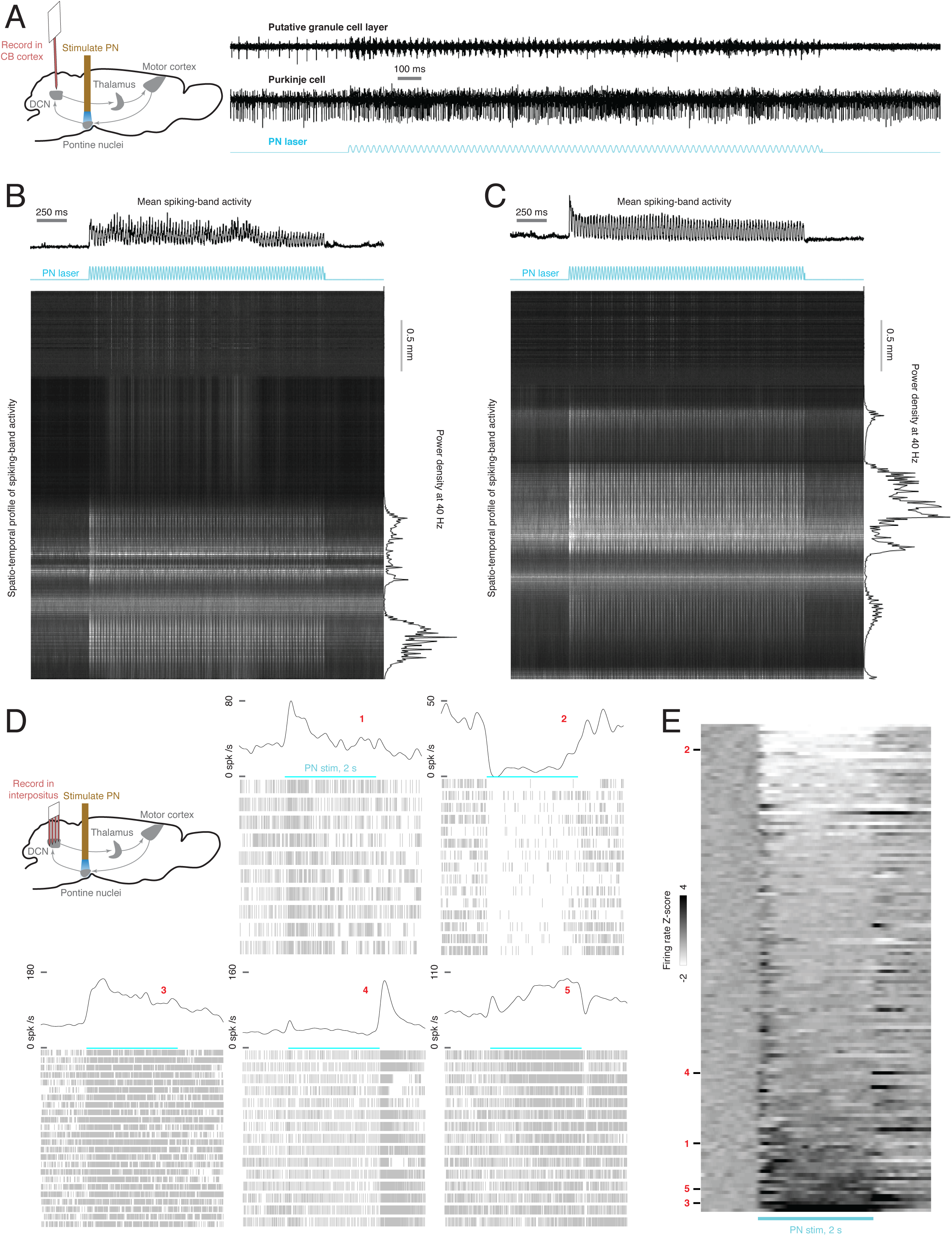
Effect of optogenetic stimulation of the PN on the cerebellum. (A), Left: schematic of experiment. ChR2 expression was driven in Slc17a7-Cre mice by viral injection in the PN, and an optical fiber was implanted in the PN. Multiunit activity was recorded in cerebellar cortex with a 384-channel Neuropixels probe as a two-second, 40 Hz sinusoidal laser stimulus was delivered to the PN. Right: raw data examples from a recording site in the Purkinje cell layer and a site in the granule cell layer. (B) Spatio-temporal profile of the cerebellar cortical response to PN stimulation. The heatmap shows raw data from each channel of the probe, filtered and rectified, during the PN stimulus (see Methods). The upper trace shows multi-unit activity over time, averaged across all channels. The right trace shows the power density at 40 Hz for each channel, as a function of depth. (C) Same as (B); data from a different mouse. (D) Upper left: experimental schematic. A two-second, 40 Hz laser stimulus was applied to the PN while activity was recorded in the deep cerebellar nuclei (DCN). Lower and right: laser-aligned spike times and firing rates for five example DCN neurons. (E) Firing rate z-scores for all DCN neurons (n = 139), aligned to PN stimulus onset.

**Figure 4:**
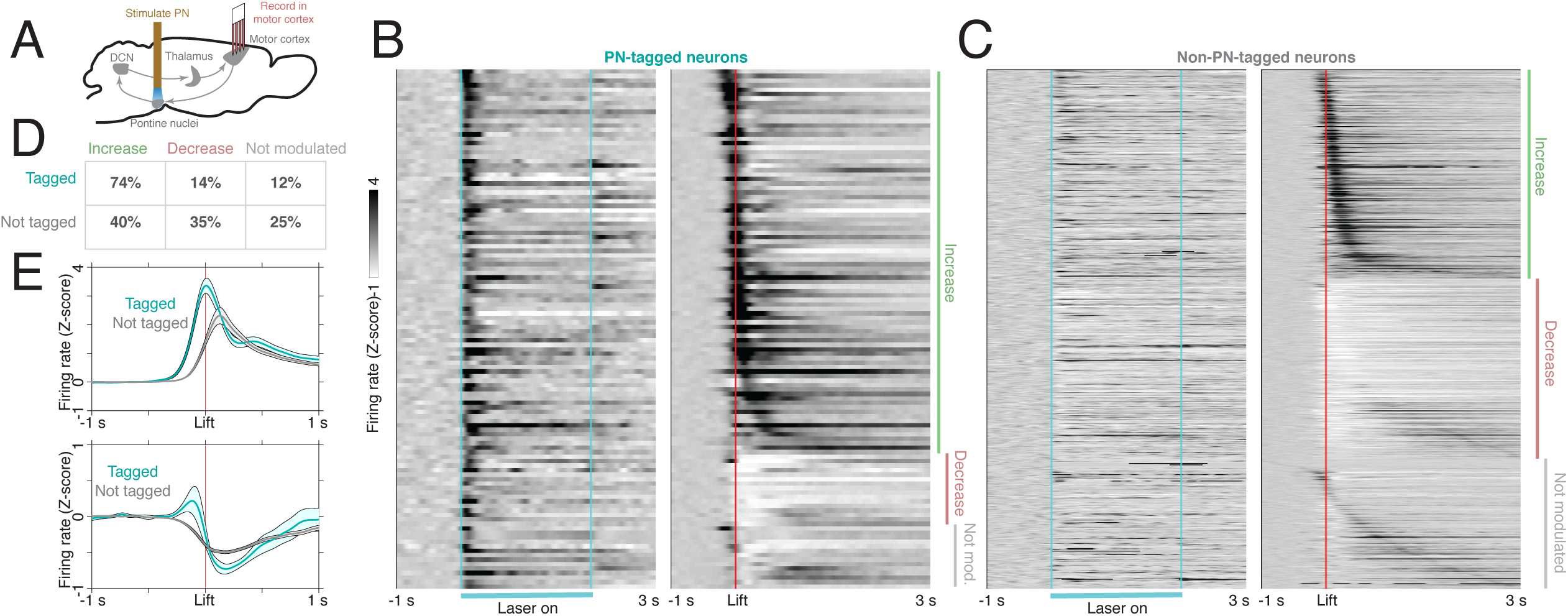
Motor cortical neurons receiving feedback from the ponto-cerebellar system have distinct functional properties during reaching. (A) Experimental schematic. A 40 Hz, 2 s stimulus was delivered to PN neurons to identify motor cortical cells receiving PN feedback. Then, the activity of PN-tagged and non-PN-tagged neurons was compared during normal reaching. (B) Laser-aligned and movement-aligned neural activity for PN-tagged cortical neurons (n = 116/848); n = 26 sessions, n = 8 mice. Many neurons had increased activity around lift, and a few had decreased activity. (C): Laser-aligned and movement-aligned neural activity for non-PN-tagged cortical neurons (n = 732). Approximately equal numbers of neurons had firing rate increases and decreases around lift. (D) Table showing the percentages of tagged and non-tagged neurons with increases, decreases, or no change in activity around lift. (E) Upper: average firing rate Z-scores for tagged and non-tagged neurons with firing rate increases around lift. The average activity for the tagged group increased earlier than the activity for the non-tagged group. Lower: average firing rate Z-scores for tagged and non-tagged neurons with firing rate decreases around lift.

### Motor cortical neurons receiving ponto-cerebellar feedback have distinct functional profiles

The binary nature of cortical responses to PN stimulation allowed us to classify cortical neurons into a group that was tagged by stimulation and a group that was not tagged. We examined the firing patterns of PN-tagged and non-PN-tagged cortical neurons during the reaching behavior, by comparing pre- and post-lift spike counts (see Methods). While both tagged and non-tagged neurons exhibited lift-locked increases and decreases, firing rate increases were much more common among tagged neurons (Fig. 4B-D; chi-square test, p = 8.1e-11). Furthermore, among the cells with firing rate increases, the tagged neurons increased their activity earlier and had a larger peak than the non-tagged neurons (Fig. 4E, upper), consistent with previous findings in primates using electrical stimulation of cerebellar output (Nashef et al., 2018). The difference in movement-related activity between tagged and untagged neurons might occur because cerebello-cortical inputs are driving early responses in motor cortex. Alternatively, inputs to cortex that convey cerebellar signals may, during development or learning, preferentially target cortical neurons that are active early in the movement.

### Optogenetic perturbation of the PN perturbs hand kinematics and impairs reach-to-grab performance

How does stimulation of the PN influence movement execution? In order to address this, we had animals perform the reaching task, and trials with laser stimulation of the PN were interleaved with control trials. PN stimulation produced a range of effects on reach kinematics in different mice (Supplemental Fig. 3). First, in some cases, the reach began normally, but the hand overshot the target, and the animal grabbed at a location past the pellet (Fig. 5A-C, left and center panels). A difference in the position of the hand at grab was observed in at least one direction in 15/45 sessions from 9/15 mice, and the largest differences occurred in the forward direction, corresponding to overreaching (Fig. 5D). Second, the delay from lift to grab increased relative to control, reflecting a slower reach (Fig. 5A-C, right panels; Fig. 5E); this occurred in 27/45 sessions from 11/15 mice. Third, in a few cases (5/45 sessions from 4/15 mice), stimulation reduced the probability of movement initiation (Fig. 5F). Fourth, stimulation reduced the rate of success on the first reach attempt in 14/45 sessions from 8/15 mice (Fig. 5G; one session showed the opposite effect). Fifth, the dispersion in the position of the hand at grab was higher on laser trials in 17/45 sessions from 8/15 mice. At least one effect was observed in 32/45 sessions from 14/15 mice. Thus, while PN stimulation disrupted skilled reaching, the specific kinematic effects differed across animals. This finding is consistent with clinical observations of cerebellar ataxia patients (Klockgether, 2000).

**Figure 5:**
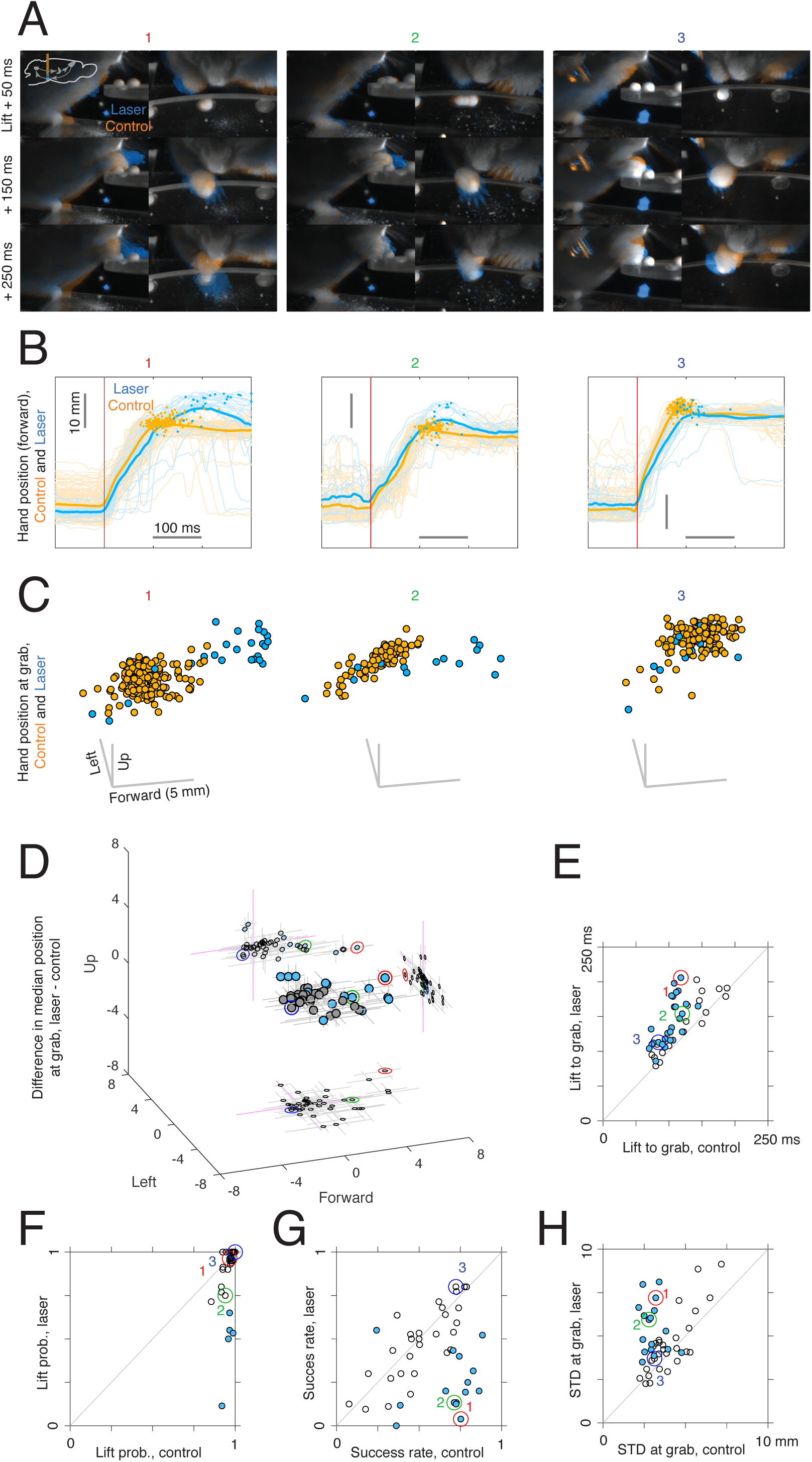
Effects of PN stimulation on reaching. Mice expressing ChR2 in the PN were implanted with a fiber to allow 40 Hz, 2 s stimulation of PN neurons. Neural activity in the motor cortex, cerebellar nuclei, or both was also recorded in these sessions. (A) Video frames for control and PN laser stimulation trials for sessions from three mice. Frames from the front and side cameras are shown at offsets of 50, 150, and 250 ms from lift, averaged across trials (see Methods). Orange indicates higher image intensity on control trials, and blue indicates higher intensity on laser trials. For the first two mice (left panel), this visualization shows that the animals reach farther forward on laser trials than control trials. (B) Hand trajectories for the three example mice in (A). Dots indicate the position of the hand at grab. (C) Three-dimensional position of the hand at the time of grab on control and laser trials for each of the three mice in (A-B). Each point corresponds to a single trial. (D) Difference in median hand position at grab between laser and control trials. Each point corresponds to a single experimental session. Filled blue dots correspond to sessions in which a difference between control and laser was found in at least one direction (two-sided rank sum test for each direction, q < .05). Lines indicate bootstrapped 95% confidence intervals. (E) Median delay from lift to grab, laser vs control; filled dots indicate sessions where a difference between laser and control was detected (two-sided rank-sum test, q < .05). (F) Probability of initiating a reach, laser vs control; filled dots represent sessions where a difference between control and laser was detected (chi-square test, q < .05). (G) Success rate on first reach attempt for laser vs control; filled dots indicate a difference between laser and control (chi-square test, q < .05). (H) Standard deviation (summed across forward, left, and up directions) in the position of the hand at grab for laser vs control trials. Each point corresponds to one experimental session, and filled dots represent sessions where a difference between control and laser was detected in at least one direction (two-sample F-test for equal variances, two-sided, q < .05).

### Effects of PN stimulation on the cerebellar nuclei and motor cortex during reaching

How does PN stimulation affect neural dynamics in the DCN and motor cortex during reaching? We recorded from the DCN during reaching on control trials, in which the animal reached to the pellet following a cue, and laser trials, when PN stimulation was delivered throughout the trial (Fig. 6A, lower inset; n = 139 neurons from n = 12 sessions in n = 8 mice). On control trials, DCN neurons were strongly modulated around movement (Fig. 6A, orange traces; Fig. 6B; n = 60 neurons with firing rate increases, and n = 70 neurons with firing rate decreases), consistent with previous results in primates (Fortier et al., 1989, 1993) and mice (Becker and Person, 2019). Lift-aligned activity on laser trials was typically similar to the control pattern (Fig. 6A, left and center), but in some cases, the stimulation abolished the normal pattern of activity (Fig. 6A, right). In order to quantify the similarity between neural activity on control and laser trials, we first computed the correlation between the firing rates of all neurons on control trials and all neurons on laser trials at each peri-lift time point. This correlation became positive shortly before movement onset and remained high throughout the movement: the cells that had higher firing rates at a given time on control trials also tended to have higher firing rates on laser trials at that time (Fig. 6C, upper). Next, for each neuron, we computed the correlation between firing rates on control and laser trials over the entire peri-lift window; this provides an estimate of how similar the control and laser patterns are for individual neurons. These correlations tended to be positive and large (positive for 131/139 neurons, negative for 5/139 neurons; permutation test with large-sample approximation, q < .05). Thus, PN stimulation did not erase the patterns of activity that occurred during normal movement, but instead modulated these patterns (Fig. 6D). The laser-induced firing rate changes during the task were correlated with the changes predicted by neurons’ response to the laser alone (Supplemental Fig. 4A-E), except in a short period around movement onset (Supplemental Fig. 4E, upper).

**Figure 6:**
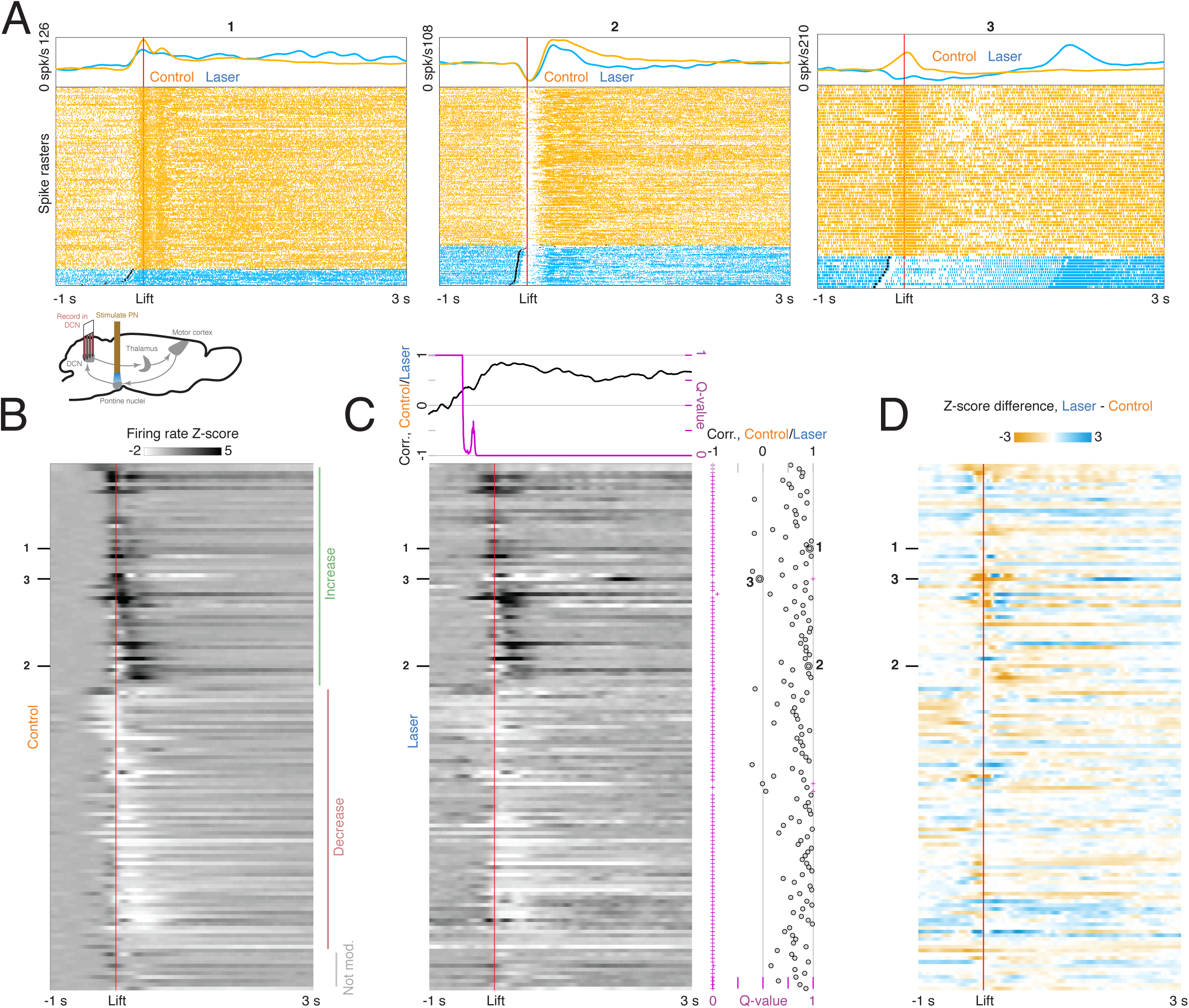
Effect of PN stimulation on neural activity in the deep cerebellar nuclei (DCN) during reaching. (A) Lower inset: experimental schematic. Animals performed the reaching task as neural activity in the deep cerebellar nuclei was recorded. On some trials, PN neurons were activated with a 40 Hz, 2 s laser stimulus. Top panels: lift-aligned spike times and firing rates on control and laser trials for three DCN neurons. (B) Heatmap displaying lift-aligned average firing rate z-scores on control trials for all DCN neurons (n = 139). (C) Heatmap displaying lift-aligned average firing rate z-scores for laser trials. Upper inset: the black line shows the rank correlation (Spearman’s ρ) between firing rates of the 139 neurons on control and laser trials at each time point. The magenta inset shows the q-value against the null hypothesis that the correlation is zero. Right inset: black circles show the rank correlation (Spearman’s ρ) between the control and laser firing rates over time for each neuron. Magenta crosses show the q-value against the null hypothesis that the correlation of control and laser values over time is zero. (D) Difference between lift-aligned z-scores for control and laser trials. Orange regions indicate neurons and time points in which the firing rate is higher on control trials, and blue regions indicate points in which the firing rate is higher on laser trials.

The modulation of DCN activity by PN stimulation might perturb reaching kinematics through descending routes to the spinal cord, or by altering activity in motor cortex (Fig. 1B). We therefore recorded from motor cortical ensembles (n = 1157 neurons from n = 38 sessions in n = 10 mice) during reaching on trials with PN stimulation and on control trials (Fig. 7A, lower inset). Similarly to DCN neurons, cells in motor cortex were strongly modulated during movement on control trials, and often exhibited small differences on laser trials (Fig. 7A). Firing rates on control and laser trials were positively correlated over time for most cells (Fig. 7B-C; positive correlation for 965 neurons and negative correlation for 47 neurons; permutation test with large-sample approximation, q < .05). However, as with DCN activity, cortical activity during the task was modulated by PN stimulation (Fig. 7D). Consistent with the observation that cortical responses to laser-only stimulation were mainly transient increases (Supplemental Fig. 5A), the effect of PN stimulation during the task was correlated with the effect predicted from the laser-only responses immediately before movement onset (Supplemental Fig. 5B-E). Other differences between activity on control and laser trials, which included decreases in activity and could not be accounted for by laser-only responses (Supplemental Fig. 5D-E), were observed later during the movement.

**Figure 7:**
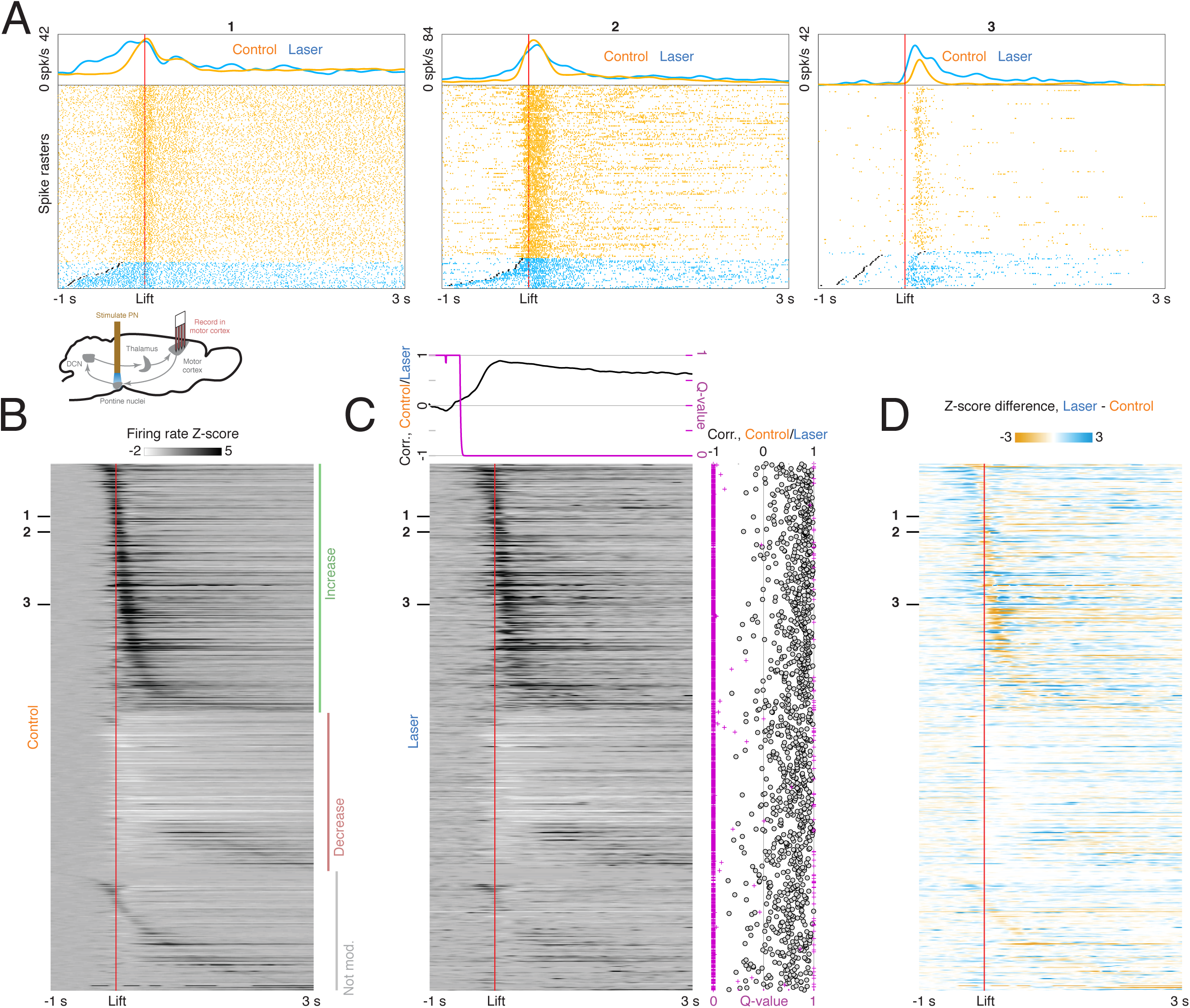
Effect of PN stimulation on neural activity in motor cortex during reaching. (A) Lower inset: experimental schematic. Animals performed the reaching task as neural activity in motor cortex was recorded. On some trials, PN neurons were activated with a 40 Hz, 2 s laser stimulus. Top panels: lift-aligned spike times and firing rates on control and laser trials for three motor cortex neurons. (B) Heatmap displaying lift-aligned average firing rate z-scores on control trials for all motor cortex neurons (n = 1157). (C) Heatmap displaying lift-aligned average firing rate z-scores for laser trials. Upper inset: the black line shows the rank correlation (Spearman’s ρ) between firing rates of the 1157 neurons on control and laser trials at each time point. The magenta inset shows the q-value against the null hypothesis that the correlation is zero. Right inset: black circles show the rank correlation (Spearman’s ρ) between the control and laser firing rates over time for each neuron. Magenta crosses show the q-value against the null hypothesis that the correlation of control and laser values over time is zero. (D) Difference between lift-aligned z-scores for control and laser trials. Orange regions indicate neurons and time points in which the firing rate is higher on control trials, and blue regions indicate points in which the firing rate is higher on laser trials.

### Decoding of hand velocity from neural activity on control and PN stimulation trials

Are the differences in neural activity during the movement consistent with the kinematic differences between control and PN stimulation trials? In order to address this question, we designed linear filters to decode 3D hand velocities from population activity recorded during control trials in motor cortex and DCN (n=38 sessions with cortex recordings, n=10 sessions with DCN recordings, of which n=5 had simultaneous recordings in both areas; Figure 8A, Supp. Figure 6A-B, Methods). We then compared the trajectories decoded from neural activity in trials with PN stimulation with those decoded in a test set of control trials not used for training the decoder. We found that the decoded trajectories were different between the two trial types (Figure 8B-C, center columns), confirming that the population activity in cortex and DCN was altered by the perturbation of PN in directions relevant to behavior. The differences in decoded velocities between the two trial types qualitatively matched the differences observed in behavior (Figure 8B-C, left vs center columns) with statistically significant correlations in each direction for the cortical decoding (Figure 8B, right columns) and in the lateral direction for DCN decoding (Figure 8C, right columns). The fact that the neural decoders better reproduced the observed velocities in control test trials than laser trials (Supp. Figure 6C-D) could be due to actual differences in how the neural signals are transformed into behavior (e.g. via other compensatory pathways active when PN are perturbed). However, these differences could also be related to generalization error since the decoder was only trained on control trials. An alternative procedure in which the decoder was trained on a balanced set of control and laser trials yielded similar performance on both trial types (Supp. Figure 6E-F, Methods), suggesting that in both regions there are dimensions of neural activity that can explain the behavior equally well in perturbed and control trials. Overall, these results show that the changes in the activity in cortex and DCN induced by the PN stimulation during movement are consistent with the behavioral effects of the perturbation.

**Figure 8:**
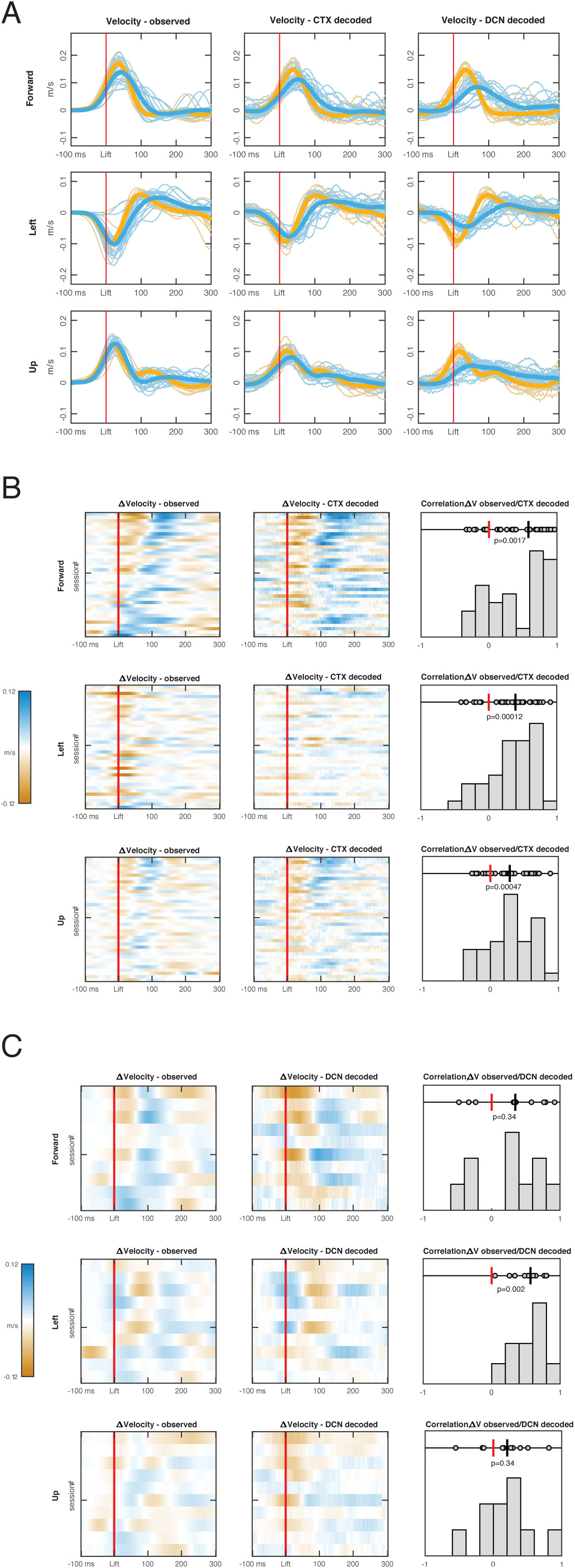
Hand velocity decoding from neural activity in cortex (CTX) and deep cerebellar nuclei (DCN). (A) Observed (left column) vs decoded (center CTX, right DCN) velocity trajectories in each direction for a session with simultaneous recordings. Blue lines represent all trials with PN perturbation (“laser” trials) and their mean (thicker line). Orange lines represent all “test control” trials, not used for training the decoder, and their mean. (B) Observed vs decoded mean differences in velocity between laser and control test trials, for each of the sessions with CTX recordings (n = 38). In the heatmaps (left observed, center decoded) rows are sessions, ordered based on the observed mean velocity difference in the forward direction in the 100ms following lift. The horizontal tick denotes the session shown in (A). Right panels show the distribution of rank correlations (Spearman’s ρ) between the observed and decoded differences in velocity over time (i.e. between corresponding rows of the heatmaps). Within each panel, individual correlations and their median (black line) are shown on top, with p-value of the two-sided sign rank test against the null hypothesis of zero median (red line), and histogram of correlation values across sessions is shown at the bottom. (C) Same as (B), for each of the sessions with DCN recordings (n=10).

## Discussion

One way to test the function of a neural loop is to identify a node that projects to only one other node within the loop and no structures outside it. If such a node can be identified and experimentally manipulated, the effects of this manipulation on the activity patterns of downstream and upstream nodes can be related to behavioral effects. We investigated the contribution of the cortico-cerebellar loop during behavior by stimulating the PN, a feedforward node in this loop which has no intrinsic recurrent connections and projects only to the cerebellum. This approach allowed us to disrupt signals from the forebrain into the cerebellum; by contrast, a direct perturbation of motor cortex would have influenced not only the cerebellum, but also spinal circuits and movement, via corticospinal, cortico-reticulospinal, and cortico-rubrospinal routes, complicating the interpretation of our results. In particular, our targeted approach enabled us to study how a disruption of descending cortico-cerebellar signals impacted activity not only in the cerebellum, but also across the full loop in motor cortex, which itself projects back to the perturbation site in the PN.

We stimulated the pontocerebellar pathway in the absence of reaching to characterize the signal transformation properties of the cerebello-cortical loop. We were surprised to find that PN stimulation had a diverse range of effects on DCN neurons. If the PN influenced the DCN primarily through inhibitory Purkinje cells, one would expect sustained activation of the PN to tonically suppress the DCN. We found, however, that many DCN neurons exhibited minimal steady-state firing rate changes during laser-only trials, or even tonic increases. One possible explanation is that excitation through direct PN-DCN collaterals overrode Purkinje cell inhibition. Another possibility is that inhibitory interneurons in the cerebellar cortex partially cancelled the PN excitation at the level of granule cells or Purkinje cells. Responses to PN stimulation in motor cortex differed from those in the DCN in several key respects. First, cortical responses were smaller than those in the DCN, suggesting that the cerebello-cortical pathway attenuates PN signals, rather than amplifying them. Second, cortical responses were much more transient than those in the DCN, suggesting that the cerebello-cortical pathway can filter out steady-state firing rate changes in the DCN. Third, pontine stimulation only led to increases in cortical firing rates, in contrast with the diverse and often multiphasic responses in the DCN, suggesting that the cerebello-cortical pathway might, under certain conditions, rectify cerebellar output. These observations show how neural activity can be transformed across the cortico-cerebellar loop, and may constrain future models of the dynamics in this loop and their role in motor control.

Stimulation of the PN in a cued reaching task did not usually prevent initiation of the movement, but impaired motor skill by altering a range of kinematic features, including reach-to-grasp time, endpoint position, and endpoint variance. This range of effects resembles the dysmetria observed in spinocerebellar ataxia patients, but does not replicate other deficits, such as intention tremor. Effects resembling the weakness or paralysis that can occur following a stroke that damages motor cortex or the blocking of movement following optogenetic silencing of murine motor cortex (Galiñanes et al., 2018; Guo et al., 2015; Miri et al., 2017; Sauerbrei et al., 2020) were rarely observed. Although our perturbation only disrupted one channel of cerebellar inputs, its effects on behavior were consistent with those obtained in previous studies by manipulation of cerebellar output. In primates, high-frequency electrical stimulation of the superior cerebellar peduncle increased the tortuosity and variability of reaches (Nashef et al., 2019), consistent with our finding that PN stimulation increases endpoint variance. Previous work in mice has shown that optogenetic stimulation of the interpositus nucleus decreases outward reach velocity, and silencing the same region increases the outward velocity (Becker and Person, 2019). Taken together with previous work, our findings suggest that the cortico-cerebellar loop fine-tunes arm kinematics during skilled reaching.

Firing patterns in DCN during the behavior were affected by pontine perturbation. These changes were subtle for most neurons, but were not random; rather, they predicted the observed changes in kinematics, as revealed by decoding of hand velocity from neural activity. Perturbation of the PN could influence behavior through routes that descend from the cerebellum to the spinal cord through the red nucleus and reticular formation, or through feedback effects on motor cortex, which is critical for generating the reach (Guo et al., 2015; Sauerbrei et al., 2020). Our finding that PN stimulation modifies cortical activity during reaching, and that the modified patterns are consistent with the observed kinematic effects suggests that this stimulation alters arm movement, at least in part, through its influence on motor cortex. Thus, while the full output of motor cortex drives reaching, the cortico-cerebellar loop appears to fine-tune neural dynamics and movement kinematics.

Neural activity for reaches during PN stimulation largely recapitulated the patterns observed during normal reaching. Why might the neural dynamics be robust to disruption of one of the largest inputs to the intermediate and lateral cerebellum? Both DCN and cortex receive inputs from multiple loops that do not include pontine nuclei. Since our pontocerebellar perturbations do not directly affect these other inputs, it is likely that activity in these other loops override the aberrant PN activity and entrain relatively normal activity in the DCN and cortex. For DCN, potential inputs include sensory streams, such as those from external cuneate nucleus (Huang et al., 2013), or motor inputs, such as those from the red nucleus (Beitzel et al., 2017). These inputs may act by directly influencing the DCN or by driving near normal Purkinje cell output from the cerebellar regions that they target. Cortical patterns may be largely preserved due to the robustness of DCN activity, although other non-cerebellar inputs to cortex may contribute to compensation. These findings suggest that future work exploring the production of skilled behaviors will need to consider how dynamics in the cortico-cerebellar system integrate with those of other loops between the forebrain, hindbrain, spinal cord, and musculoskeletal plant.

## Methods

### Transgenic mouse lines

Slc17a7-Cre mice were generated by the Janelia Research Campus Gene Targeting and Transgenics Facility, and Pde1c-Cre cryopreserved sperm (line IT106) were obtained from GENSAT (Chip Gerfen; National Institutes of Health, Poolesville, MD) and rederived by the Janelia Research Campus Gene Targeting and Transgenics Facility. For the PN stimulation experiments, we used Slc17a7-Cre mice (n = 15 for behavior and cortex / DCN electrophysiology, n = 2 for cerebellar cortex electrophysiology). For the electrophysiological recordings in the PN, we obtained mice (n = 6) expressing ChR2 in pyramidal tract neurons by crossing the Cre driver line (Tg(Sim1-Cre)KJ18Gsat, The Jackson Laboratory) to a Cre-dependent ChR2 reporter mouse, Ai32 (Rosa-CAG-LSL-ChR2(H134R)-EYFP-WPRE, The Jackson Laboratory).

### Stereotaxic surgeries

All mice (2-15 months old) were anesthetized with 2% isoflurane and placed on a heating pad in a stereotactic frame (Kopf Instruments, Tujunga, CA). Mice were administered 0.1 mg/kg of subcutaneous buprenorphine at the time of surgery, and 5 mg/kg of subcutaneous Ketoprofen at the end of surgery, followed by an additional dose once a day for two days. The scalp was sanitized with three alternating rounds of iodine surgical scrub and alcohol, a portion of the scalp was removed, the skull was cleaned, and a custom-made headpost was affixed with UV-curing cement (OptiBond, Kerr; Calibra Universal Self-Adhesive Resin, Dentsply Sirona) or dental acrylic (RelyX Unicem, 3M). For viral injections into the PN, craniotomies were made using a dental drill over the PN (3.90 mm posterior to bregma, 0.4 mm lateral). Injections were made on the left (contra-limb) side. Injection pipettes were made from glass capillaries pulled on a Sutter P-97 (Sutter, Novato, CA). Viruses were loaded using a Narishige pneumatic injector (MO-10, Tokyo, Japan) and injected into the PN at 5.9, 5.6, and 5.3 mm below the dural surface. Injection volumes were 100 nl each. After each injection, at least two minutes elapsed before proceeding to the next depth, and the pipette was withdrawn five minutes after the final injection. For the ChR2 PN stimulation experiments, Slc17a7-Cre animals (n = 15 for behavior and cortex / DCN electrophysiology, n = 2 for cerebellar cortex electrophysiology) were injected with AAV-2/1-CAG-flex-ChR2-TdTomato and implanted with tapered optical fibers (Pisanello et al., 2017) in the left PN (Optogenix, Lecce, Italy; NA 0.39, taper angle 6.9°, active length ∼1mm, core diameter 200µm, core+cladding 225µm). The pattern of ChR2 expression and the placement of the fiber were assessed in postmortem histology (Supplemental Fig. 2). For the PN optogenetic perturbation experiments, a craniotomy targeting forelimb motor cortex (bregma +0.5, left 1.7 mm), the deep cerebellar nuclei (bregma -6.3, right 1.8 mm), or the cerebellar cortex (bregma -7.0 mm, right 2.5 mm) was sealed with silicone elastomer (Kwik-Sil, WPI).

Craniotomies in motor cortex and the cerebellar nuclei were performed in ten and eight mice, respectively, with three of these mice receiving craniotomies in both regions. Recordings in the cerebellar cortex were performed in two mice. Following surgery, injections of ketoprofen (5 mg/Kg) and buprenorphine (0.1 mg/Kg; Henry Schein Animal Health, Melville, NY) were administered subcutaneously. If animals exhibited signs of pain or distress following surgery, additional doses of either ketoprofen or buprenorphine were administered, as directed by veterinary staff. All procedures were performed in accordance with protocols approved by the Institutional Animal Care and Use Committee (IACUC) of the Janelia Research Campus.

### Histology

Mice were anesthetized and euthanized with isoflurane until no breathing was visually detected for ∼30 s. Mice were then transcardially perfused with 1x PBS (Fisher Scientific Inc., 20ml), followed by 4% paraformaldehyde (formalin, 20ml, Fisher Scientific, Inc). Brain tissue was removed, incubated in formalin for 24-72 hrs, then rinsed and stored in 1x PBS. Brains were sectioned at 70 μm thickness on a vibratome (Leica VT-1200S). Tissue sections were placed on glass slides and coverslipped with Vectashield DAPI HardSet Mounting Medium (Vectorlabs). Images were collected at ∼5x magnification using a fluorescent light stereo microscope (Olympus MVX10 or Nikon Eclipse Ti inverted widefield).

### Reach-to-grab task

As described previously (Guo et al., 2015; Sauerbrei et al., 2020), mice undergoing behavioral training were acclimated to head restraint and food restricted to 80-90% of original body weight by limiting food intake to 2-3 g/day. Otherwise, mice had ad libitum food. Animals were monitored daily by veterinary staff, according to IACUC guidelines. During training, animals learned to reach for pellets of food with the right paw following an acoustic cue. In most sessions, the pellet was delivered with a rotating table (Janelia Experimental Technology), as described previously (Guo et al., 2015; Sauerbrei et al., 2020). In n = 7 sessions with DCN recording, food was instead delivered on an automated vertical post (Janelia Experimental Technology). Two high-speed cameras (Point Grey Flea3) with manual iris and focus lenses (Tokina 6-15 mm f/1.4, or Tamron 13VM1040ASIR 10-40mm, f/1.4) were placed in front and to the right of the animal. A custom-made infrared LED light source was mounted behind each camera. Video was recorded at 500Hz using BIAS acquisition software (IO Rodeo, available at https://bitbucket.org/iorodeo/bias). The cameras, acoustic cue and table, and laser were controlled using Wavesurfer software (Adam Taylor, Janelia Scientific Computing; http://wavesurfer.janelia.org/) and a custom Arduino controller (Peter Polidoro, Janelia Experimental Technology).

### Video analysis

The position of the hand was tracked using the APT software package (https://github.com/kristinbranson/APT), developed by the Branson Lab at Janelia, as described previously (Guo et al., 2015; Sauerbrei et al., 2020). The position of the hand was manually annotated for training frames, a tracker was created using the cascaded pose regression (Dollár et al., 2010) or DeepLabCut algorithm (Mathis et al., 2018), and the tracker was applied to all movies in each dataset. The three-dimensional position of the hand was triangulated by performing a stereo calibration of the pair of cameras using the Caltech Camera Calibration Toolbox for Matlab (http://www.vision.caltech.edu/bouguetj/calib_doc/). The timing of behavioral waypoints, including lift and grab, was estimated with the Janelia Automatic Animal Behavior Annotator (https://github.com/kristinbranson/JAABA), as described previously. Lift was defined as initial separation between paw and perch. Hand-open was defined as fingers beginning to separate from palm. Grab was defined as paw moving downwards as digits closed. Supination was defined as wrist rotation >90° upwards. At-mouth was defined as the frame in which the paw was within 1 pixel of mouth. Chew was defined as mastication of pellet visible in mouth. Each behavior classifier inputs a short sequence of frames from both the front- and side-view videos, and outputs a prediction of whether the mouse is performing the given behavior or not in the center frame (note that a different classifier is trained for each behavior and each mouse). The classifier used Histogram of Oriented Gradient (HOG) and Histogram of Optical Flow (HOF) features, general-purpose features that represent the directions and magnitudes of edge and motion vectors. Following the initial classification, post-processing was performed in which the per-frame classification results were smoothed by filling short gaps, and spurious short detection bouts were removed. We further filtered the detected events by considering only event sequences in which a lift, hand-open, and grab were detected in order. Using the JAABA interface, the user manually checked for and corrected classifier errors and retrained the classifier, if necessary. Trials were regarded as “single-reach successes” if the first lift-hand-open-grab sequence resulted in the animal grabbing the pellet, bringing it to the mouth, and chewing. Trials were regarded as “multi-reach successes” if the animal missed on the first attempt, but subsequently grabbed the pellet, brought it to the mouth, and chewed.

### Electrophysiological recordings from the PN

On the day prior to recording, a craniotomy was made over the PN (A/P: -3.5-4.0mm, M/L: 0.2-0.5mm, D/V: 5.5-6.0mm) in Sim1-Cre X Ai32 mice (n = 6) and sealed with silicone elastomer (Kwik-Sil, WPI). On the recording day, the animal was head-fixed, a fiber coupled to a 473 nm laser (LuxX 473-80, Omicron Laserage) was placed over forelimb motor cortex, and a 384-channel Neuropixels 3A probe (https://www.neuropixels.org/) coated in a fluorescent dye (CM-DiI, DiI, DiO, Thermofisher Scientific; JF-669, Tocris) was slowly lowered into the brain. As the probe approached the PN, layer 5 motor cortical neurons were activated with a 10 Hz train of 5 ms pulses (2-10 mW), and the depth profile of evoked spiking was examined online. When it was determined that the bottom ∼1 mm of the probe was in the PN, the craniotomy and recording probe were sealed with Kwik-Sil, and after 15 min the reaching task was initiated. Neuropixels data and timestamps for the camera and laser were acquired with a custom FPGA-based system. Electrophysiological data were spike sorted using JRClust (https://github.com/JaneliaSciComp/JRCLUST). For offline assessment of the depth profile of evoked spiking, the data for each channel were blanked from 0-6 ms following pulse onset to remove the optical artifact, high-pass filtered with a cutoff of 650 Hz, full-wave rectified, averaged over all pulses, and smoothed over time with a σ = 333 μs Gaussian kernel. Next, the data were spatially smoothed across the probe using a σ = 30 μm Gaussian kernel, and the depth profile averaged between a temporal offset of 6-10 ms from pulse onset was plotted (Fig. 2B, red curve). Units on the bottom 1 mm of the probe were included for analysis; this is the region that the histology and depth profiles suggested to be in the PN. Firing rates were visualized by smoothing the cue-aligned spike trains using a σ = 50 ms Gaussian kernel, Z-scoring based on the mean and standard deviation of the firing rate from cue - 1000 ms to cue - 200 ms, and averaging across trials (Fig. 1E.) For the long laser stimuli (Fig. 2G, upper), spike counts were compared between the 2 s laser-on period and the 2 s pre-laser period using a two-sided sign rank test. Multiple comparisons were corrected for by controlling the false discovery rate at .05 (Benjamini and Hochberg, 1995). Firing rates were classified as changing during the laser if the corrected p-value q < .05.

### Generalized linear model (GLM) for cue and reach responses in the PN

In order to disentangle the effects of the cue and movement on PN activity, we fit a generalized linear model (GLM) to spiking activity for each PN neuron (Park et al., 2014). In this model, spikes are generated by an inhomogeneous Poisson process with intensity given by

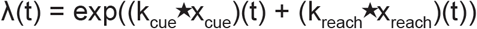

where x_cue_ and x_reach_ are delta functions centered at cue and lift times, k_cue_ and k_reach_ are linear filters describing the time-varying effects of each event type, and ⋆ denotes convolution. The filters k_cue_ and k_reach_ were constructed using half-cosine basis functions. Eight cosine bumps were evenly spaced across an 800 ms window starting at the event time. The model was fit by maximizing the log posterior with ridge regularization on the weights using the neuroGLM package for Matlab: https://github.com/pillowlab/neuroGLM. We assessed the goodness-of-fit (R^2^) by regressing the observed firing rates (Gaussian smoothing with σ = 50 ms) on the estimated firing rate λ(t). We visualized the estimated effects of the cue and reach on PN activity by plotting exp(k_cue_) and exp(k_reach_) for the neurons with R^2^ > 0.1 (Fig. 2F, three upper left panels and lower two heatmaps). To assess the effects of cue and reach, we computed the Euclidean norm of k_cue_ and k_reach_ and compared their distributions (Fig. 2F, upper right).

### PN-stimulation-evoked multi-unit activity in the cerebellum

In order to determine whether the effects of optogenetic stimulation of the PN propagated into cerebellar cortex, we stimulated ChR2-expressing PN neurons with an implanted optical fiber, and inserted a 384-channel Neuropixels probe into cerebellar cortex (bregma -7.0 mm, right 2.5 mm, depth 2.5 mm, with the probe four degrees from vertical, so that the tip of the probe was anterior to the point at which it entered the brain). We then delivered light in a 40 Hz sinusoidal pattern to the PN (mean power 3 mW). Because we were able to obtain very few stable, well-isolated neurons in the cerebellar cortex, we instead examined the spatiotemporal profile of multi-unit activity, which likely reflects the combination of signals from mossy fiber terminals (including those from stimulated PN neurons), granule cells, Purkinje cells, and interneurons. Data from each site on the Neuropixels probe were high-pass filtered with a cutoff of 650 Hz, full-wave rectified, and smoothed over time with a Gaussian kernel (σ = 333 μs). The resulting traces were spatially smoothed across the probe (Gaussian kernel, σ = 30 μm), and the resulting spatio-temporal profile was visualized as a heatmap, with the channel-averaged signal and power density at 40 Hz for each site displayed as insets (Fig. 3B-C).

### Optogenetic stimulation of the PN

Selective expression of ChR2 was induced in the PN by injecting Slc17a7-Cre mice (n = 15) with AAV-2/1-CAG-flex-ChR2-TdTomato in the left PN and implanting a tapered optical fiber over the injection site, as described above. Animals were trained on the reach-to-grab task, and craniotomies over either forelimb motor cortex, the DCN (interpositus nucleus), or both were performed before the first day of recording. During the recording session, an optical coupler attached to a 473 nm laser (LuxX 473-80, Omicron Laserage) was attached to the connection on the top of the skull leading to the tapered fiber. During the experiment, three types of trials were administered: (1) control trials, in which a cue was given and the food pellet delivered, (2) laser-only trials, in which a 2 s, 10 Hz, 20 Hz, or 40 Hz sinusoidal laser stimulus was given with no cue, and (3) laser-cue trials, in which the sinusoidal laser stimulation was given either synchronously with the cue, 200 ms before the cue, or 300 ms before the cue. In most sessions (n = 38/45), 40 Hz stimulation was used, but in some (n = 7/45), 10 Hz or 20 Hz stimulation was applied because pilot behavioral experiments on those mice suggested that lower-frequency stimulation might have larger effects on behavior. The laser power at the tip of the coupler was measured at 8-20 mW, but there was likely a significant power drop between the coupler and the fiber in the brain. To directly visualize the effect of PN stimulation in the raw video frames, we compiled the video frames at offsets of 50 ms, 150 ms, and 250 ms from lift, and computed the 90th percentile of image intensity for each pixel on control trials and laser trials, resulting in one frame template for control and laser at each temporal offset. Then, we used the control and laser templates as pixelwise weights for orange (RGB = [1 .5 0]) and blue (RGB = [0 .5 1]), respectively, and added these weighted values together to produce the final images (Fig. 5A). Hand trajectories were plotted as a function of time (Fig. 5B), and the position of the hand at the time of grab was extracted for control and laser plots and compared by plotting (Fig. 5C). In order to summarize the effect of PN stimulation for each session, we used five kinematic and task variables, as follows. To correct for multiple comparisons across the 45 sessions, the Benjamini-Hochberg correction was applied to the p-values from the statistical tests for each measure, and we rejected the corresponding null hypotheses if the corrected p-value q < .05.

1. Difference in the median position of the hand at grab on laser and control trials (Fig. 5D; two-sided rank-sum test for each spatial direction, q < .05).
2. Median time from lift to grab on control and laser trials (Fig. 5E; two-sided rank-sum test, q < .05).
3. Probability of initiating a lift on control and laser trials (Fig. 5F; chi-square test, q < .05).
4. Success rate on the first reach attempt on control and laser trials (Fig. 5G; chi-square test, q < .05).
5. Standard deviation in the position of the hand at grab for laser and control trials (Fig. 5H shows the sum of the standard deviation across the three spatial directions; for each direction, a two-sample, two-sided F-test for equal variances was performed, q < .05, and dots are blue for sessions in which the null hypothesis was rejected for at least one direction).

We were unable to make reliable comparisons of cue-to-reach reaction time on laser and control trials across sessions, due to three technical issues. First, in some sessions in which we delivered the pellet on a vertical post, the reaction times were negative on control trials, because the animal could hear the post raising up and reached before the acoustic cue was given. Second, we imposed a delay from the laser onset to the cue in several sessions, because we had previously observed an apparent tendency for animals to reach at the laser onset, and wanted to verify that animals were not using the laser as a cue. Third, PN stimulation blocked movement initiation in some sessions, which resulted in an uninterpretable reaction time on laser trials for those sessions. Because these issues complicated the interpretation of cue-to-reach reaction time, we did not include it as a variable of interest.

### Electrophysiological recordings in the DCN and motor cortex

At the start of the optogenetic stimulation experiments (n = 45 sessions, n = 15 mice), four-shank, 64-channel probes (Janelia Experimental Technology) were lowered to approximately 0.9 mm below the surface of forelimb motor cortex, 2.5 mm below the cerebellar surface, or both. In total, we obtained recordings from 1157 cortical neurons across 38 sessions in ten mice and recordings from 139 DCN neurons across 12 sessions in five mice. For five of these sessions in three mice, both areas were recorded simultaneously. For the DCN (interpositus nucleus), the probe was lowered until large, high-frequency spikes were visible and complex spikes (characteristic of Purkinje cells in the cerebellar cortex) were no longer present. The 16 recording sites on each shank had a range of zero to 320 μm from the tip of the shank. Spike sorting was performed with Kilosort2 (Stringer et al., 2019) (https://github.com/MouseLand/Kilosort2) or JRClust (https://github.com/JaneliaSciComp/JRCLUST). In order to assess whether cortical neurons were modulated during movement and by PN stimulation, we compared paired spike counts for each trial immediately before and after lift or laser onset (using laser-only trials) with a sign-rank test (two-sided for lift, one-sided for laser). The pre- and post-lift windows were (−1000, -500) and (−50, 450), and the pre- and post-laser windows were (−150, 0) and (0, 150) ms. A Benjamini-Hochberg correction for multiple comparisons was applied, and neurons were classified as modulated by the lift or laser if the corrected p-value q < .05. Event-aligned firing rates were visualized by smoothing the spike trains with a σ = 50 ms Gaussian kernel, Z-scoring based on the mean and standard deviation of the firing rate within a window of (−1000, -400) ms of lift and (−1000, 0) of laser, and averaging across trials. For DCN neurons, the same procedure was used to characterize movement-aligned responses. Because the laser-only responses of DCN neurons were extremely variable, exhibiting a mixture of transient and tonic increases and decreases, we could not classify them into tagged and untagged groups, as we did for cortical neurons. For both cortex and DCN neurons, we computed the Spearman correlation in firing rates on control and laser trials at each time point across all neurons (Fig. 6C and 7C, upper insets), and the Spearman correlation across all time points for each individual neuron (Fig. 6C and 7C, right insets). In order to determine whether the firing rate changes following laser-only stimulation were related to firing rate changes induced by laser stimulation during the movement, we predicted what effect the laser would have had on neural activity if it induced the same response during reaching as during laser-only stimulation. To do that, we took the peri-laser-only firing rate for each neuron, z-scored it using the pre-lift mean and standard deviation, offset it at the observed lift-to-laser delays for each trial in the corresponding session, and averaged across trials (Supp. Fig. 4E, 5E, lower heatmaps). We then computed the Spearman correlation between these predictions and the actual laser - control firing rate z-score differences across all neurons at each time point (Supp. Fig. 4D, 5D). The correlation values over time, along with the q-values for the test against the null hypothesis of zero correlation, are plotted in Supp. Fig. 4E, 5E (upper insets).

### Neural decoding

We designed linear filters to decode 3D hand velocities from neural activity recorded in motor cortex (CTX) and the deep cerebellar nuclei (DCN), to assess whether the observed behavioral differences between control trials and trials with PN perturbation (“laser trials”) were related to corresponding differences in neural activity. To guarantee that decoding results for laser trials reflect movements performed with PN perturbed, we excluded laser trials in which the lift-to-grab sequence was not initiated and completed while the laser was turned on or in which the lift occurred less than 300 ms before the end of the laser period (since the decoding window extends up to 300 ms after lift). Two sessions with DCN recordings had less than two laser trials satisfying these criteria, so they were not included in the decoding analysis. For sessions in which the pellet was delivered on the rotating table, we also excluded atypical control trials in which the lift-to-grab sequence was not completely included within the two seconds following the acoustic cue. For sessions in which the pellet was delivered on a vertical post, we used a similar criterion but also allowed the lifts to occur up to 0.5 s before the cue because the motor moving the pellet started earlier than the acoustic cue, and some animals effectively cued their movement to the motor. Following our prior work (Sauerbrei et al., 2020), the decoders use multi-unit neural activity preprocessed as follows: (a) counts of detected spikes are smoothed with a Gaussian kernel with σ = 25 ms, (b) these smoothed firing rates are z-scored with respect to the activity at rest (1.5s window preceding the start of each trial) and channels with mean absolute z-scores greater than 100 during movement are excluded, (c) these z-scores are processed with PCA, with the principal components computed from the lift-aligned trial-averaged activity in control trials (window -100 ms to 300 ms around lift). At any instant of time, the decoders use the 15 most recent samples (hence up to 28 ms in the past) of PCA-reduced neural activity in CTX or DCN, to decode the hand velocity at that time. The following procedure was used to choose the number of PCA dimensions used by the decoder and the decoder coefficients in each experimental session. First, one fifth of the control trials (“control test” set) was kept aside for testing the performance of the decoder, thus enabling fair comparison with decoder performance in laser trials. On the remaining set of control trials, 4-fold cross validation was performed with an increasing number of PCA dimensions. For each fold and each number of PCA dimensions, decoder coefficients were computed by regressing observed hand velocity data against PCA-reduced neural data in three-fourth of the trials (window -100 ms to 300 ms around lift), and then used to predict the velocities in the left-out trials. Hand velocities were computed by smoothing the raw 3D hand trajectories with a Gaussian kernel (σ = 25 ms, as for neural activity) and then applying a central difference filter of order 8. For each number of PCA dimensions, we compared the average across folds of the mean squared error between predicted and observed velocities. We finally chose the minimum number of PCA dimensions that guaranteed performance within 1% of the overall minimum across all choices, and averaged the decoder coefficients for that number of PCA dimensions across the folds. The decoder thus derived was applied to the neural activity data in the control test trials and the laser trials to predict the hand velocities with and without PN perturbation and contrast these predictions with the hand velocities observed experimentally (Figure 8, Supp. Figure 6A-B). We applied the decoding analysis to all experimental sessions with cortical recordings (n = 38) and with DCN recordings (n = 10) independently, choosing in each case the region of the dataset that maximized the number of stable channels (from CTX or DCN). To summarize the decoding performance in each session, we used the coefficient of determination (R^2^) between decoded and observed hand velocities, after pooling all directions together. The velocity decoding accuracy was higher in control test trials than in laser trials in most of the CTX decoding sessions, and all of the DCN decoding sessions (Supp. Fig. 6C-D). This drop in R^2^ on laser trials could be due to actual differences in how the neural signals are transformed into behavior (e.g. via other compensatory pathways active when PN are perturbed) but could also partly reflect generalization error. In fact, the decoder was trained only on control trials, and therefore a drop in decoding performance was expected for a trial type not used for training. We investigated this possibility by repeating the decoding analysis with an alternative decoder trained on a perfectly balanced set of control and laser trials. This “balanced decoder” was similar to the original decoder, but: (a) the training set consisted of all odd laser trials and a matching number of control trials, roughly equally spaced in time within the set of all control trials, (b) PCA components were computed from the average of the lift-aligned trial-averaged activity in laser trials and the lift-aligned trial-averaged activity in control trials, (c) the number of PCA dimensions used by the decoder was fixed to 5 rather than cross-validated, (d) the decoder performance was tested on the laser and control trials not used for training the decoder. Note that while in the original decoder the test set is roughly balanced between laser and control trials, for the balanced decoder it is the training set that is exactly balanced, but the test set contains many more control trials than laser trials. When we compared the balanced decoder performance on control and laser test trials, we did not find statistically significant differences in either cortical or DCN-based decoding (Supp. Figure 6E-F), suggesting that in both regions there is a neural population that can explain the behavior during PN perturbation as well as in control trials, and the difference in performance in the original decoder may in fact be due to generalization error.

We performed additional analyses with the original decoder on the sessions with simultaneous CTX and DCN recordings (n = 5, all using the table for pellet delivery), to compare the decoding performance between the two neural populations. First, we repeated the computation of the decoder coefficients for the CTX-based and the DCN-based velocity decoders (each using the cross-validated number of PCA dimensions found in the original decoding analysis) after excluding trials that were not available in both datasets, in order to guarantee that both decoders used identical training sets. Similarly, a common set of control test and laser trials was used to compare the performance of the two decoders. Figure 8A shows the results of this additional analysis performed on one of the simultaneous sessions. The decoding performance of the CTX-based decoders was better than that of the DCN-based decoders in all of the simultaneous sessions. However, this result was confounded by the fact that the recording yield in cortex was superior to that in the DCN in all these sessions. In fact, subsampling the cortical units to match the number of recorded DCN units lowered the CTX-based decoding performance to a level comparable to that of DCN-based decoding, so we did not include the results of this analysis in the results and discussion.

## Contributions and acknowledgements

J.G. performed behavioral experiments, DCN recordings, and cortical recordings and analyzed behavior data. B.S. analyzed electrophysiology and behavior data. J.C. performed Neuropixels recordings in the PN and cerebellar cortex. M.M. developed and performed the neural decoding analyses. A.G. performed preliminary slice electrophysiology experiments and analyzed preliminary behavioral data. F.P. developed and fabricated the optical fibers for PN stimulation. K.B. developed video analysis tools and analyzed preliminary behavior data. B.S., M.M., and A.H. wrote the paper with input from all authors. A.H. supervised the project. We thank Gülşen Sürmeli, Diana Burk, and Cheng-Chiu Huang for help with pilot experiments, Adam Taylor for Wavesurfer software, Peter Polidoro for instrumentation development, Tim Harris and the Neuropixels Consortium for probe development, James Jun and Marius Pachitariu for spike sorting software, Sal DiLisio and the Janelia Vivarium for viral injections and implantation surgeries, the Janelia Histology Core for histology and imaging support, the Janelia Visitor Program for supporting pilot experiments, and Yifat Prut, Abdulraheem Nashef, Steve Edgley, Amy Bastian, Stephen Scott, Daniel Wolpert, and Brett Mensh for discussions.

## Figure captions

**Supplemental Figure 1 (related to Figure 2):**
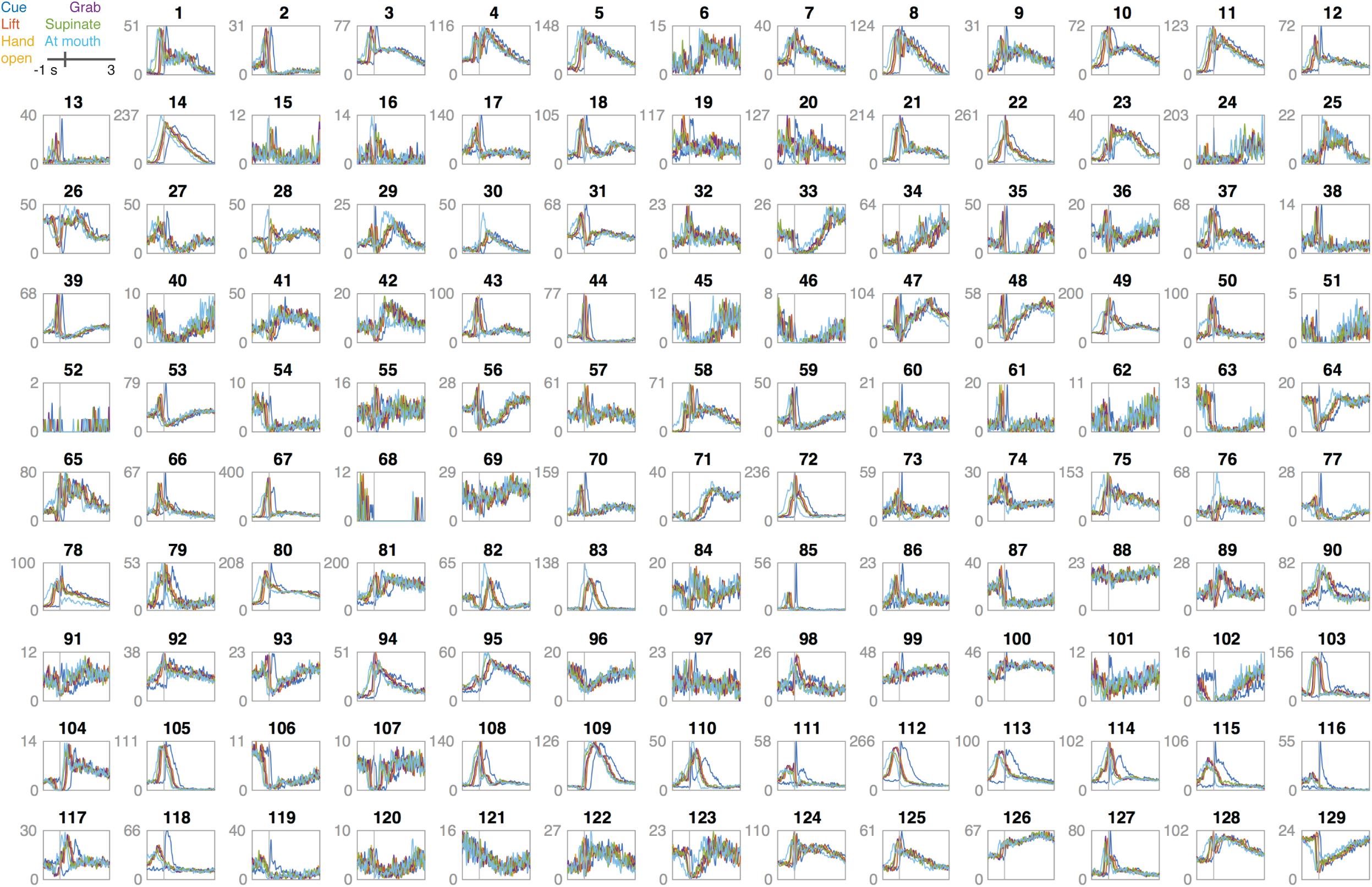
Firing rates of PN neurons aligned to different behavioral events. Each panel shows the trial-averaged firing rate of one PN neuron in a window of -1 s to +3 s of the acoustic cue, lift, hand open, grab, supination, and the arrival of the hand at the mouth.

**Supplemental Figure 2 (related to Figures 5-8):**
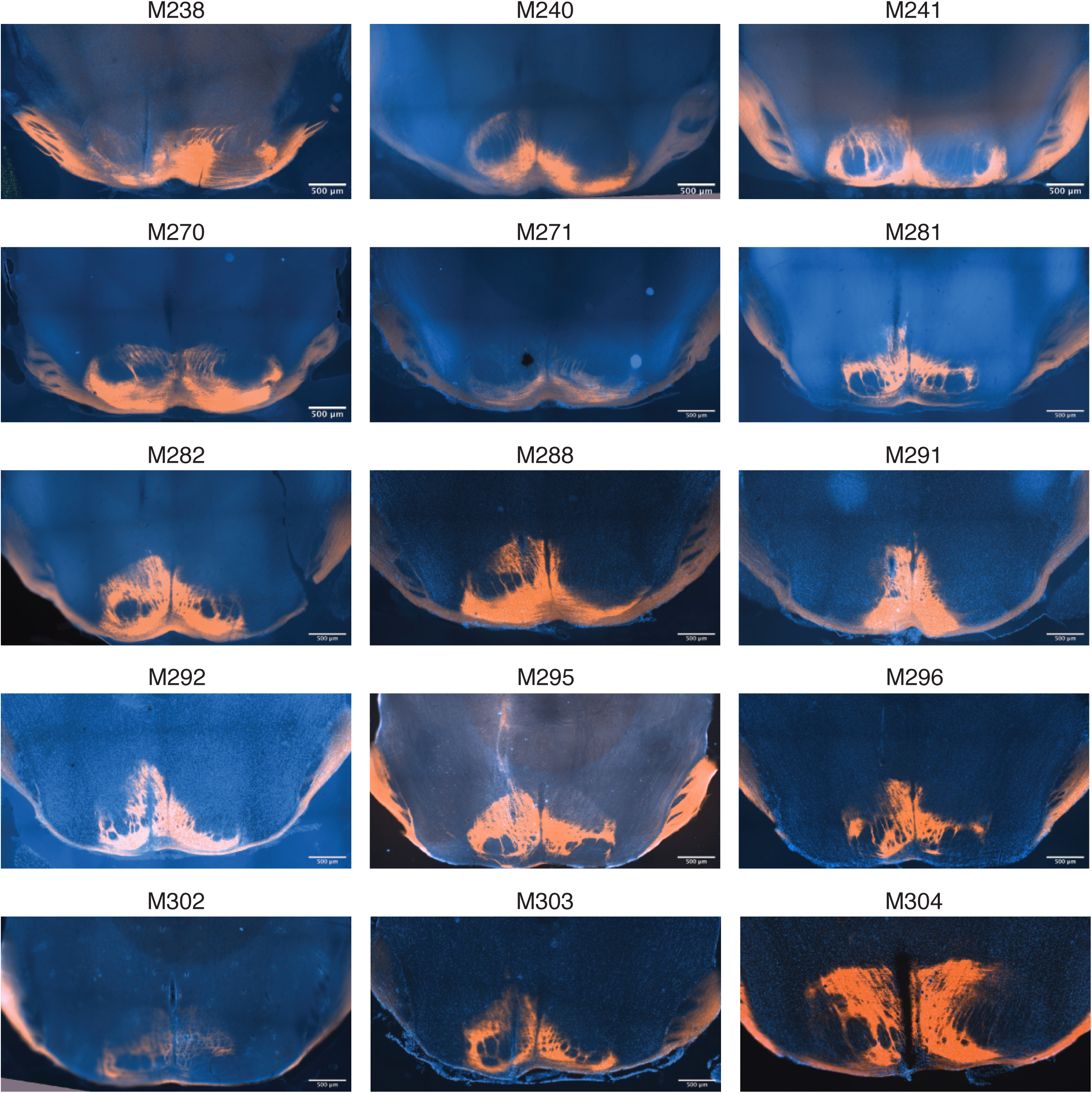
Histological sections showing the expression of ChR2 in the PN and the position of the optical fiber.

**Supplemental Figure 3 (related to Figure 5):**
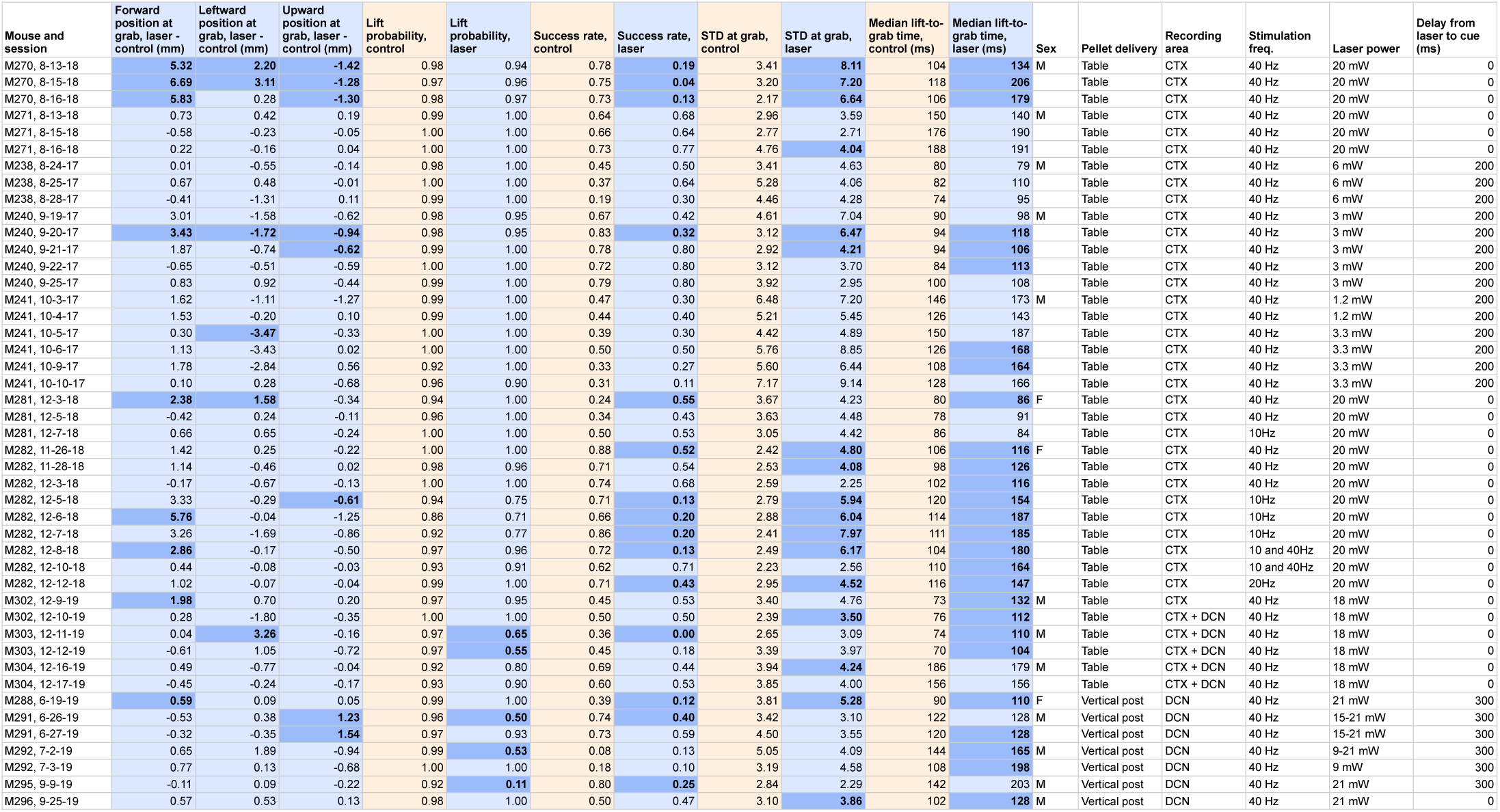
Table of effects of PN stimulation on arm kinematics and task performance. Orange columns correspond to control trials. Blue columns correspond to laser trials or average differences between laser and control trials. Dark blue entries correspond to cases in which the null hypotheses are rejected (see Methods for the statistical tests used).

**Supplemental Figure 4 (related to Figure 6):**
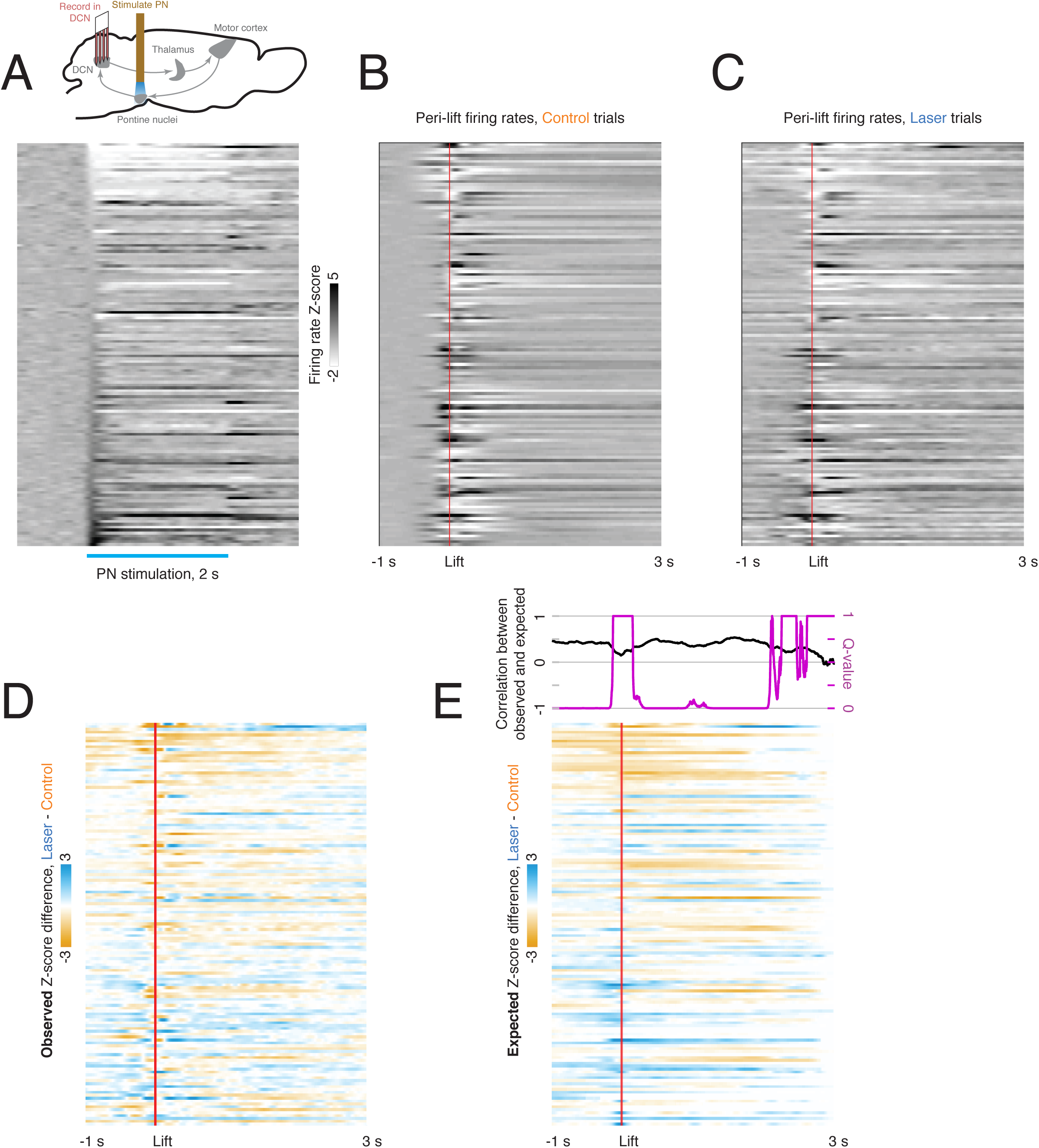
Relationship between laser-aligned and lift-aligned activity in the DCN. (A) Peri-laser firing rate z-scores for DCN neurons (n = 139). Only trials in which a laser stimulus but no cue was delivered were included. Data are sorted by the average z-scored firing rate within the first 100 ms of laser onset. (B) Peri-lift firing rates on control trials for the same neurons as in (A). (C) Peri-lift firing rates on laser + cue trials. (D) Effect of PN stimulation on neural activity in the DCN during behavior. The heatmap shows the difference in z-scored peri-lift firing rates on laser and control trials; it was obtained by subtracting the map in (A) from the map in (B). Blue indicates higher peri-lift activity on laser trials, and orange indicates higher firing rates on control trials. (E) Predicted effect of PN stimulation on neural activity during behavior based on laser-only response. For each neuron, the neural activity aligned to laser stimulation in the absence of a cue was offset at the lift-to-laser offsets in the corresponding session and averaged over trials. The resulting heatmap in the lower panel shows the lift-aligned firing rate changes expected based on the laser-only responses. The upper panel shows the correlation (Spearman’s rho) between the expected and observed effect of the laser on the neural population at each time point (black), along with the q-value for the test against the null hypothesis that this correlation is zero.

**Supplemental Figure 5 (related to Figure 7):**
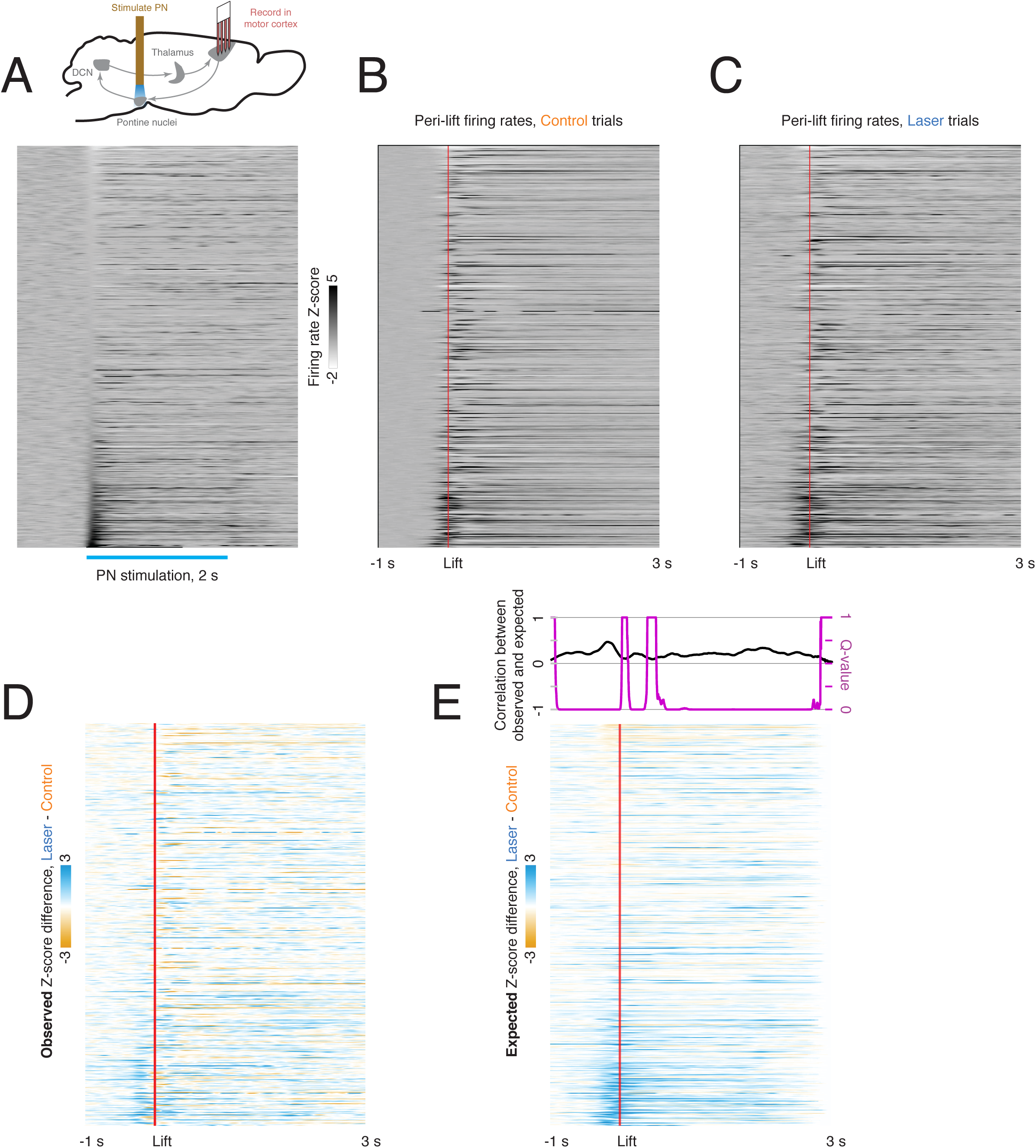
Relationship between laser-aligned and lift-aligned activity in motor cortex. (A) Peri-laser firing rate z-scores for cortical neurons (n = 839). Only trials in which a laser stimulus but no cue was delivered were included. Data are sorted by the average z-scored firing rate within the first 100 ms of laser onset. (B) Peri-lift firing rates on control trials for the same neurons as in (A). (C) Peri-lift firing rates on laser + cue trials. (D) Effect of PN stimulation on neural activity in the motor cortex during behavior. The heatmap shows the difference in z-scored peri-lift firing rates on laser and control trials; it was obtained by subtracting the map in (A) from the map in (B). Blue indicates higher peri-lift activity on laser trials, and orange indicates higher firing rates on control trials. (E) Predicted effect of PN stimulation on neural activity during behavior based on laser-only response. For each neuron, the neural activity aligned to laser stimulation in the absence of a cue was offset at the lift-to-laser offsets in the corresponding session and averaged over trials. The resulting heatmap in the lower panel shows the lift-aligned firing rate changes expected based on the laser-only responses. The upper panel shows the correlation (Spearman’s rho) between the expected and observed effect of the laser on the neural population at each time point (black), along with the q-value for the test against the null hypothesis that this correlation is zero.

**Supplemental figure 6 (related to Figure 8):**
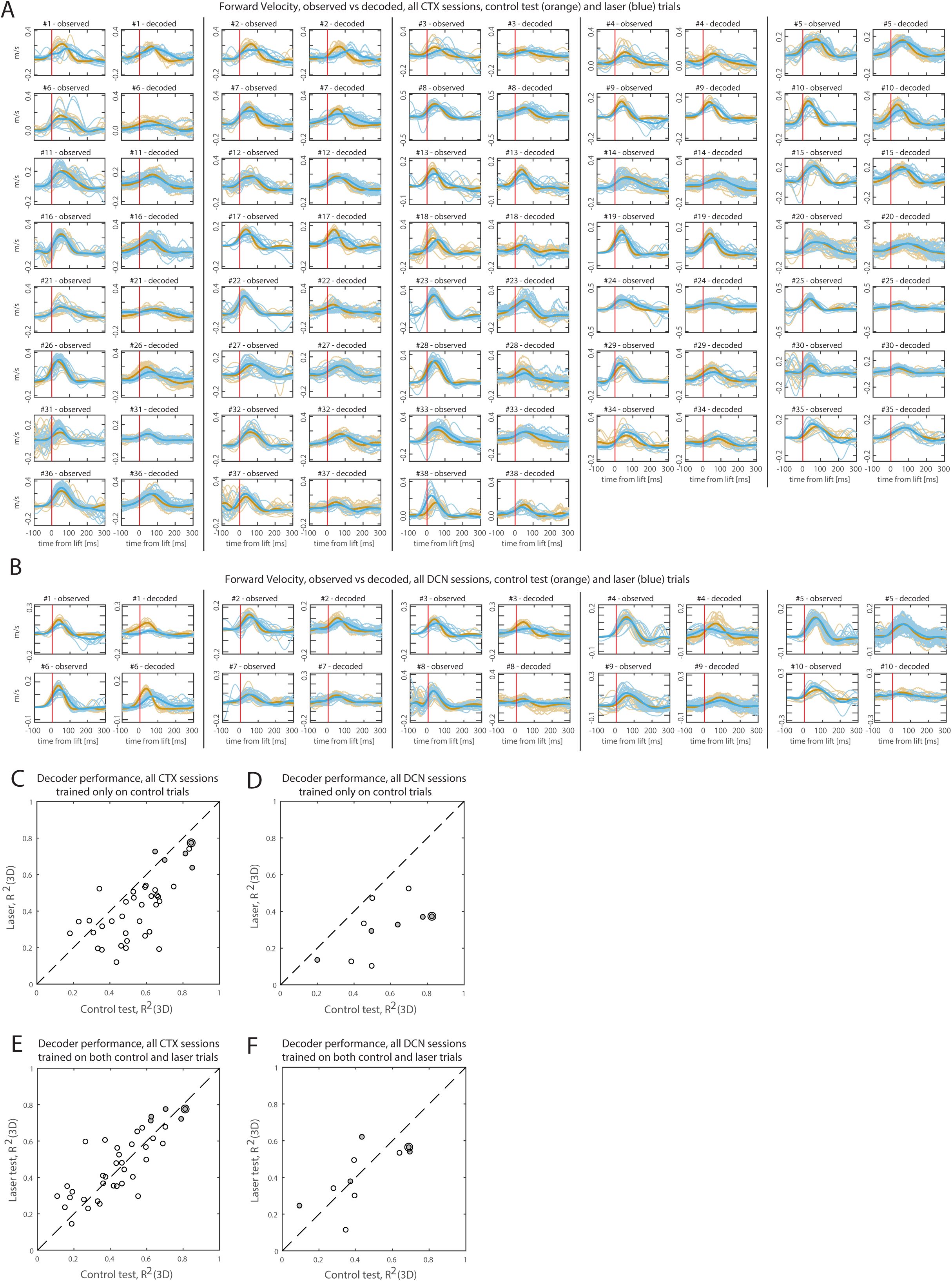
Hand velocity decoding from neural activity in motor cortex (CTX) and deep cerebellar nuclei (DCN), for all sessions with recordings in at least one of the regions (n=38 CTX sessions, n=10 DCN sessions; n=5 sessions had simultaneous recordings in both regions). (A), (B) Observed vs decoded velocity trajectories in the forward direction, for all sessions with cortical recordings (A) and DCN recordings (B). Blue lines represent all trials with PN perturbation (“laser” trials) and their mean (thicker line). Orange lines represent all “test control” trials, not used for training the decoder, and their mean. Sessions are ordered as in Fig. 8B-C. (C), (D) Performance of the decoders – which are trained only on a subset of control trials - in control test vs laser trials, for all sessions with cortical recordings (C) and DCN recordings (D). Filled circles represent sessions with simultaneous CTX & DCN recordings, and outer circle denotes session in Fig. 8A. Performance metric is the coefficient of determination (R^2^) between decoded and observed hand velocities, after pooling all directions together. For both cortex and DCN, the decoded velocities better reproduced the observed velocities in control than laser trials (in CTX, mean R^2^ for control test trials (0.55) > mean R^2^ for laser trials (0.42), p=0.002; in DCN, mean R^2^ for control test trials (0.54) > mean R^2^ for laser trials (0.30) p=0.005, two-tailed Welch t-tests). (E), (F) Performance of decoders trained on a balanced subset of control and laser trials (see Methods), for all sessions with cortical recordings (E) and DCN recordings (F). There were no significant differences in the performance of these decoders in control test trials (mean R^2^ =0.45 for CTX, 0.43 for DCN) compared with laser trials (mean R^2^ =0.47 for CTX, 0.41 for DCN).

## References

Becker, M.I., and Person, A.L. (2019). Cerebellar Control of Reach Kinematics for Endpoint Precision. Neuron 103, 335–348.e5.

Beitzel, C.S., Houck, B.D., Lewis, S.M., and Person, A.L. (2017). Rubrocerebellar Feedback Loop Isolates the Interposed Nucleus as an Independent Processor of Corollary Discharge Information in Mice. J. Neurosci. 37, 10085–10096.

Benjamini, Y., and Hochberg, Y. (1995). Controlling the False Discovery Rate: A Practical and Powerful Approach to Multiple Testing. J. R. Stat. Soc. Series B Stat. Methodol. 57, 289–300.

Berndt, A., Schoenenberger, P., Mattis, J., Tye, K.M., Deisseroth, K., Hegemann, P., and Oertner, T.G. (2011). High-efficiency channelrhodopsins for fast neuronal stimulation at low light levels. Proc. Natl. Acad. Sci. U. S. A. 108, 7595–7600.

Border, B.G., Kosinski, R.J., Azizi, S.A., and Mihailoff, G.A. (1986). Certain basilar pontine afferent systems are GABA-ergic: combined HRP and immunocytochemical studies in the rat. Brain Res. Bull. 17, 169–179.

Brodal, A., and Jansen, J. (1946). The ponto-cerebellar projection in the rabbit and cat. Experimental investigations. J. Comp. Neurol. 84, 31–118.

Brodal, P., and Bjaalie, J.G. (1992). Organization of the pontine nuclei. Neurosci. Res. 13, 83–118.

Brodal, P., Mihailoff, G., Border, B., Ottersen, O.P., and Storm-Mathisen, J. (1988). GABA-containing neurons in the pontine nuclei of rat, cat and monkey. An immunocytochemical study. Neuroscience 25, 27–45.

Brooks, V.B., Kozlovskaya, I.B., Atkin, A., Horvath, F.E., and Uno, M. (1973). Effects of cooling dentate nucleus on tracking-task performance in monkeys. J. Neurophysiol. 36, 974–995.

Cajal, S.R. (1898). Histology of the Nervous System.

Dollár, P., Welinder, P., and Perona, P. (2010). Cascaded pose regression. In 2010 IEEE Computer Society Conference on Computer Vision and Pattern Recognition, (ieeexplore.ieee.org), pp. 1078–1085.

Dow, R.S., and Moruzzi, G. (1958). The Physiology and Pathology of the Cerebellum (U of Minnesota Press).

Fogassi, L., Gallese, V., Buccino, G., Craighero, L., Fadiga, L., and Rizzolatti, G. (2001). Cortical mechanism for the visual guidance of hand grasping movements in the monkey: A reversible inactivation study. Brain 124, 571–586.

Fortier, P.A., Kalaska, J.F., and Smith, A.M. (1989). Cerebellar neuronal activity related to whole-arm reaching movements in the monkey. J. Neurophysiol. 62, 198–211.

Fortier, P.A., Smith, A.M., and Kalaska, J.F. (1993). Comparison of cerebellar and motor cortex activity during reaching: directional tuning and response variability. J. Neurophysiol. 69, 1136–1149.

Galiñanes, G.L., Bonardi, C., and Huber, D. (2018). Directional Reaching for Water as a Cortex-Dependent Behavioral Framework for Mice. Cell Rep. 22, 2767–2783.

Gao, Z., Davis, C., Thomas, A.M., Economo, M.N., Abrego, A.M., Svoboda, K., De Zeeuw, C.I., and Li, N. (2018). A cortico-cerebellar loop for motor planning. Nature 563, 113–116.

Georgopoulos, A.P., Kalaska, J.F., Caminiti, R., and Massey, J.T. (1982). On the relations between the direction of two-dimensional arm movements and cell discharge in primate motor cortex. J. Neurosci. 2, 1527–1537.

Gerfen, C.R., Paletzki, R., and Heintz, N. (2013). GENSAT BAC cre-recombinase driver lines to study the functional organization of cerebral cortical and basal ganglia circuits. Neuron 80, 1368–1383.

Guo, J.-Z., Graves, A.R., Guo, W.W., Zheng, J., Lee, A., Rodríguez-González, J., Li, N., Macklin, J.J., Phillips, J.W., Mensh, B.D., et al. (2015). Cortex commands the performance of skilled movement. Elife 4, e10774.

Huang, C.-C., Sugino, K., Shima, Y., Guo, C., Bai, S., Mensh, B.D., Nelson, S.B., and Hantman, A.W. (2013). Convergence of pontine and proprioceptive streams onto multimodal cerebellar granule cells. Elife 2, e00400.

Jenkinson, E.W., and Glickstein, M. (2000). Whiskers, barrels, and cortical efferent pathways in gap crossing by rats. J. Neurophysiol. 84, 1781–1789.

Klockgether, T. (2000). Handbook of Ataxia Disorders (CRC Press).

Krakauer, J.W., and Carmichael, S.T. (2017). Broken Movement: The Neurobiology of Motor Recovery After Stroke (MIT Press).

Lawrence, D.G., and Kuypers, H.G. (1968). The functional organization of the motor system in the monkey. I. The effects of bilateral pyramidal lesions. Brain 91, 1–14.

Leergaard, T.B., Alloway, K.D., Pham, T.A.T., Bolstad, I., Hoffer, Z.S., Pettersen, C., and Bjaalie, J.G. (2004). Three-dimensional topography of corticopontine projections from rat sensorimotor cortex: Comparisons with corticostriatal projections reveal diverse integrative organization. J. Comp. Neurol. 478, 306–322.

Legg, C.R., Mercier, B., and Glickstein, M. (1989). Corticopontine projection in the rat: the distribution of labelled cortical cells after large injections of horseradish peroxidase in the pontine nuclei. J. Comp. Neurol. 286, 427–441.

Levesque, F., Fabre-Thorpe, M., Wiesendanger, M., and Buser, P. (1986). Brachium pontis lesions in cats partly reproduce the cerebellar dysfunction of voluntary reaching movements. Behav. Brain Res. 21, 167–181.

Mathis, A., Mamidanna, P., Cury, K.M., Abe, T., Murthy, V.N., Mathis, M.W., and Bethge, M. (2018). DeepLabCut: markerless pose estimation of user-defined body parts with deep learning. Nat. Neurosci. 21, 1281–1289.

Matsunami, K. (1987). Neuronal activity in nuclei pontis and reticularis tegmenti pontis related to forelimb movements of the monkey. Neurosci. Res. 5, 140–156.

Meyer-Lohmann, J., Conrad, B., Matsunami, K., and Brooks, V.B. (1975). Effects of dentate cooling on precentral unit activity following torque pulse injections into elbow movements. Brain Research 94, 237–251.

Meyer-Lohmann, J., Hore, J., and Brooks, V.B. (1977). Cerebellar participation in generation of prompt arm movements. J. Neurophysiol. 40, 1038–1050.

Mihailoff, G.A., Lee, H., Watt, C.B., and Yates, R. (1985). Projections to the basilar pontine nuclei from face sensory and motor regions of the cerebral cortex in the rat. J. Comp. Neurol. 237, 251–263.

Miri, A., Warriner, C.L., Seely, J.S., Elsayed, G.F., Cunningham, J.P., Churchland, M.M., and Jessell, T.M. (2017). Behaviorally Selective Engagement of Short-Latency Effector Pathways by Motor Cortex. Neuron 95, 683–696.e11.

Nashef, A., Cohen, O., Israel, Z., Harel, R., and Prut, Y. (2018). Cerebellar Shaping of Motor Cortical Firing Is Correlated with Timing of Motor Actions. Cell Rep. 23, 1275–1285.

Nashef, A., Cohen, O., Harel, R., Israel, Z., and Prut, Y. (2019). Reversible Block of Cerebellar Outflow Reveals Cortical Circuitry for Motor Coordination. Cell Rep. 27, 2608–2619.e4.

Park, I.M., Meister, M.L.R., Huk, A.C., and Pillow, J.W. (2014). Encoding and decoding in parietal cortex during sensorimotor decision-making. Nat. Neurosci. 17, 1395–1403.

Passingham, R.E., Perry, V.H., and Wilkinson, F. (1983). The long-term effects of removal of sensorimotor cortex in infant and adult rhesus monkeys. Brain 106 (Pt 3), 675–705.

Pisanello, F., Mandelbaum, G., Pisanello, M., Oldenburg, I.A., Sileo, L., Markowitz, J.E., Peterson, R.E., Della Patria, A., Haynes, T.M., Emara, M.S., et al. (2017). Dynamic illumination of spatially restricted or large brain volumes via a single tapered optical fiber. Nat. Neurosci. 20, 1180–1188.

Potter, R.F., Rüegg, D.G., and Wiesendanger, M. (1978). Responses of neurones of the pontine nuclei to stimulation of the sensorimotor, visual and auditory cortex of rats. Brain Res. Bull. 3, 15–19.

Rüegg, D., and Wiesendanger, M. (1975). Corticofugal effects from sensorimotor area I and somatosensory area II on neurones of the pontine nuclei in the cat. J. Physiol. 247, 745–757.

Sauerbrei, B.A., Guo, J.-Z., Cohen, J.D., Mischiati, M., Guo, W., Kabra, M., Verma, N., Mensh, B., Branson, K., and Hantman, A.W. (2020). Cortical pattern generation during dexterous movement is input-driven. Nature 577, 386–391.

Schmahmann, J.D., Ko, R., and MacMore, J. (2004). The human basis pontis: motor syndromes and topographic organization. Brain 127, 1269–1291.

Schwarz, C., and Thier, P. (1995). Modular organization of the pontine nuclei: dendritic fields of identified pontine projection neurons in the rat respect the borders of cortical afferent fields. J. Neurosci. 15, 3475–3489.

Stein, J.F., and Glickstein, M. (1992). Role of the cerebellum in visual guidance of movement. Physiol. Rev. 72, 967–1017.

Stringer, C., Pachitariu, M., Steinmetz, N., Reddy, C.B., Carandini, M., and Harris, K.D. (2019). Spontaneous behaviors drive multidimensional, brainwide activity. Science 364, 255.

Tziridis, K., Dicke, P.W., and Thier, P. (2009). The role of the monkey dorsal pontine nuclei in goal-directed eye and hand movements. J. Neurosci. 29, 6154–6166.

Wagner, M.J., Kim, T.H., Kadmon, J., Nguyen, N.D., Ganguli, S., Schnitzer, M.J., and Luo, L. (2019). Shared Cortex-Cerebellum Dynamics in the Execution and Learning of a Motor Task. Cell 177, 669–682.e24.

Wiesendanger, R., and Wiesendanger, M. (1982). The corticopontine system in the rat. II. The projection pattern. J. Comp. Neurol. 208, 227–238.

